# IL-17 signalling is critical for controlling subcutaneous adipose tissue dynamics and parasite burden during chronic Trypanosoma brucei infection

**DOI:** 10.1101/2022.09.23.509158

**Authors:** Matthew C. Sinton, Praveena Chandrasegaran, Paul Capewell, Anneli Cooper, Alex Girard, John Ogunsola, Georgia Perona-Wright, Dieudonné M. Ngoyi, Nono Kuispond, Bruno Bucheton, Mamadou Camara, Shingo Kajimura, Cécile Bénézech, Annette MacLeod, Juan F. Quintana

## Abstract

In the skin, *Trypanosoma brucei* colonises the subcutaneous white adipose tissue (scWAT) and harbours a pool of parasites that are proposed to be competent for forward transmission. The interaction between parasites, adipose tissue, and the local immune system is likely to drive the adipose tissue wasting and weight loss observed in cattle and humans infected with *T. brucei*. However, mechanistically, the events leading to scWAT wasting are not fully understood. Here, using several complementary approaches, including mass cytometry by time of flight, bulk and single cell transcriptomics, and *in vivo* genetic models, we found that *T. brucei* infection drives local expansion of several IL-17A-producing cells in the murine WAT, including T_H_17 and Vγ6^+^ T cells. We also found that global IL-17 deficiency, or mice lacking IL-17 receptor expression exclusively in adipocytes, were protected from infection-induced WAT wasting and weight loss. Unexpectedly, we found that abrogation of IL-17 signalling in adipocytes results in a significant accumulation of *Dpp4^+^ Pi16^+^* interstitial preadipocytes and a higher burden of extravascular parasites in the WAT, highlighting a critical role for IL-17 signalling in controlling preadipocyte fate, scWAT tissue dynamics, and local parasite burden. Taken together, our study highlights the central role of adipocyte IL-17 signalling in controlling WAT responses to infection, suggesting that adipocytes are a critical coordinator of the tissue dynamics and immune responses to *T. brucei* infection.

## Introduction

*Trypanosoma brucei* is an extracellular protozoan parasite that infects humans and livestock, causing Human African Trypanosomiasis (HAT, or sleeping sickness) and Animal African Trypanosomiasis (AAT, or nagana), respectively^1^. Both HAT and AAT are prevalent in sub-Saharan regions of the African continent, where they impose a significant socio-economic burden, and are fatal if left untreated^2^. Chronic infections in both humans and non-primate mammalian hosts, such as domestic cattle, lead to significant weight loss, a phenomenon that remains largely unstudied^3^. Upon infection, trypanosomes proliferate and migrate into tissues throughout the body, where they persist and form extravascular reservoirs in virtually every organ^4^. One major consequence of infection is weight loss, typically coupled with a reduction in white adipose tissue (WAT) mass^5^. Previous studies have elegantly demonstrated that mice lose weight during *T. brucei* infection and that this is associated with loss of gonadal white adipose tissue (gWAT) mass^6^. The largest cell volume within the gWAT is comprised of adipocytes, but this tissue is also enriched with multiple immune cell types, including macrophages, neutrophils, T helper 1 (T_H_1) cells, effector CD8^+^ cytotoxic T cells, and B cells^7–9^, as well as forming a reservoir for *T. brucei*^6^, and different factors released from these immune cells influence gWAT function under normal physiological conditions. For example, tumour necrosis factor (TNF) and interleukin-17A (IL-17A) have been shown to regulate adipose tissue structure and function, by limiting tissue expansion^7^, and inhibiting adipogenesis^9^, respectively. Furthermore, IL-17A, and signalling through the IL-17C receptor, have been shown to induce thermogenesis in white and brown adipose tissue, respectively^10, 11^, and activation of thermogenesis, following challenges such as cold exposure, leads to increased energy expenditure^12^. *T. cruzi* and *T. congloense* infections, the causative agents of Chagas disease and AAT, respectively, are also associated with elevation of IL-17A^13, 14^, which is important for controlling resistance to infection^14, 15^. Together this may suggest that loss of adipose tissue mass and subsequent weight loss may be driven by local immune responses, the parasites, or both. However, it remains unclear how the immune response to *T. brucei* infection influences adipose tissue structure and function.

Previous studies from our lab identified the skin as another reservoir for *T. brucei* and highlighted the presence of parasites in the adjacent subcutaneous white adipose tissue (scWAT) of infected patients^16^. Furthermore, we recently demonstrated a critical role of γδ T cells (in particular IL-17-producing Vγ6^+^ T cells) in the control of local skin responses to *T. brucei* infection^17^. However, we were unable to profile the scWAT in detail. Due to its proximity to the skin, the scWAT may also prove to be an important reservoir for *T. brucei*, giving the parasites access to a plentiful nutrient supply, whilst simultaneously increasing the chances of onward transmission. Therefore, we focused on understanding the impact of infection on the structure and function of the inguinal white adipose tissue (iWAT) in mice, which is typically used to model scWAT in humans^18^. Like gWAT, the iWAT acts as an energy reservoir under homeostatic conditions and modulates systemic metabolism and appetite^19^, as well as immune responses^20^. Here we present data demonstrating that *T. brucei* infection is associated with a broad immune response in the iWAT, including the expansion of IL-17A-producing T_H_17 and Vγ6^+^ T cells. Furthermore, we demonstrate that global genetic ablation of IL-17A/F or targeted deletion of adipocyte IL-17A receptor (*Il17ra*; essential for mediating IL-17A/F signalling^21^) prevents or limits weight loss and iWAT wasting. Targeted deletion of adipocyte *Il17ra* also results in an accumulation of small adipocytes in both naïve and infected mice, with a concomitant accumulation of *Dpp4^+^ Pi16^+^* interstitial preadipocytes. Interestingly, targeted deletion of adipocyte *Il17ra* also resulted in an increased burden of extravascular parasites in the iWAT. These results provide novel insights into the role of IL-17 signalling as a regulator of adipose tissue structure, function, and dynamics during infection, placing adipocyte-mediated responses at the core of the local immune responses in the iWAT. Furthermore, these findings support the utility of *T. brucei* infection models for interrogating the role that IL-17 signalling plays in controlling adipocyte fate and differentiation, as well as local and systemic energy balance, in the context of infection.

## Results

### *T. brucei* infection results in reduced iWAT mass and impaired adipose tissue function in a sex-dependent manner

In both humans and livestock, trypanosome infections are known to cause weight loss, and this has been recapitulated in male mouse models of infection^6^. However, direct comparisons have not been made between male and female mice to assess whether infection induces weight loss in a sexually dimorphic manner. To test this, we infected age-matched male and female C57BL/6 mice for a period of 25 days. We first wanted to determine that mice were successfully infected and whether there were differences between the levels of circulating parasites between sexes. Parasitaemia measurements followed a characteristic pattern, with no significant differences between sexes (**Figure 1A**). There were also no significant differences in the clinical scores of the mice (**Figure 1B**). Strikingly, during the course of infection, infected male mice lost significant amounts of bodyweight, whereas there was no significant difference between the weights of naïve and infected female mice (**Figure 1C** and **Figure 1D**). Spleen mass increased similarly in both male and female mice (**Supplementary Figure S1**), suggesting that changes in spleen mass during infection do not explain the differences in the bodyweight of males and females.

**Figure 1.**
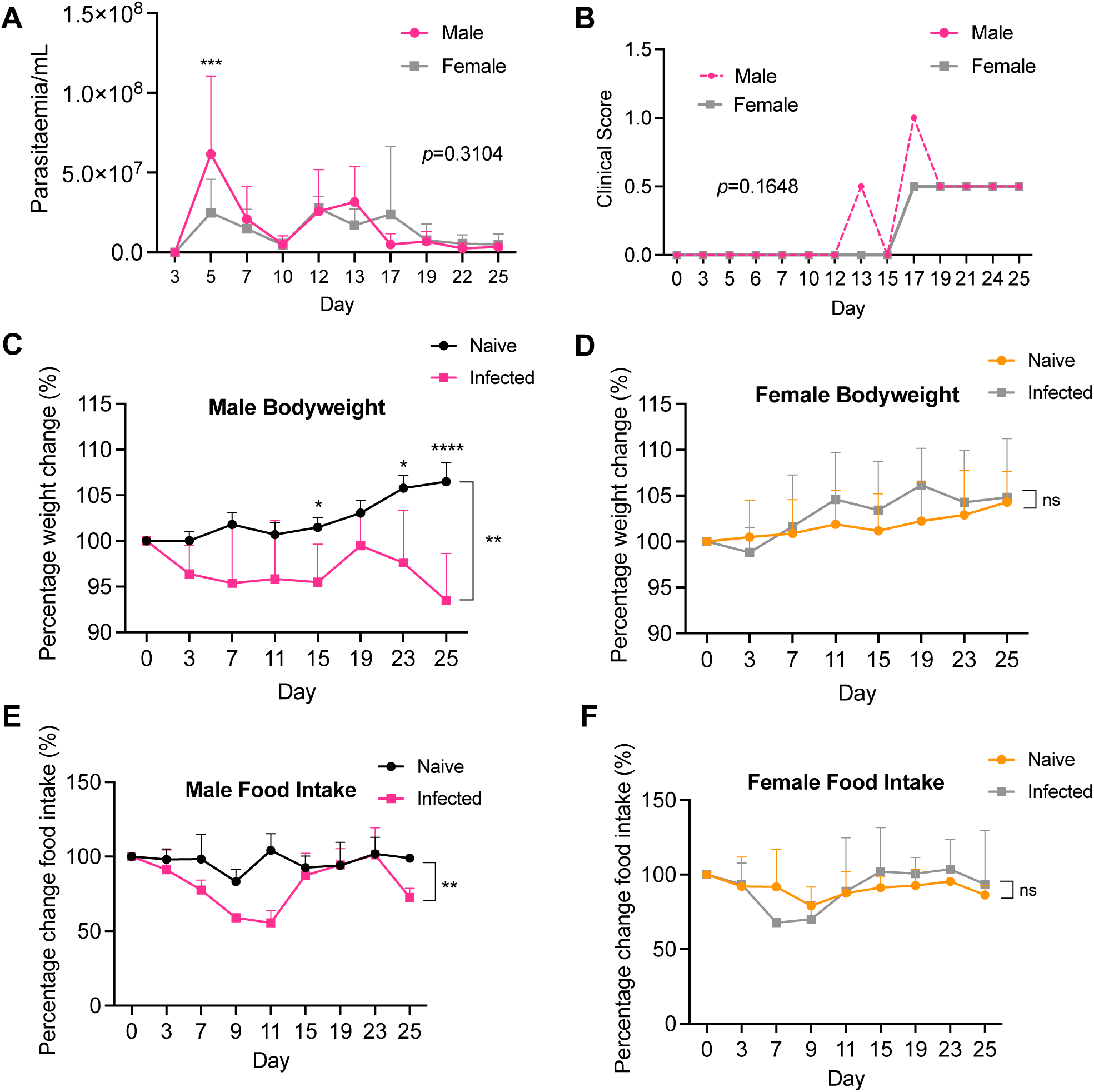
Male mice lose weight during infection with *Trypanosoma brucei*, which is associated with alterations in feeding behaviour. **(A)** Number of parasites per mL of blood, measured using phase microscopy and the rapid “matching” method^53^. **(B)** Clinical scores of infected male and female mice. **(C)** Percentage changes in body weight of male and **(D)** female mice over the course of infection. Percentage changes in food intake in male **(E)** and female **(F)** mice. Each data point represents 2 cages (*n*=3-4 mice per cage). Time series data were analysed using two-way repeated measures ANOVA with Sidak post-hoc testing and are expressed as mean ±SD. \**p*<0.05, \*\**p*<0.01, \*\*\**p*<0.001, ns = non-significant.

Weight loss may be explained as a consequence of adipose tissue wasting^22^, as a consequence of changes in feeding behaviour^23^, or both. To understand this in more detail, we measured gross food intake as a proxy for feeding behaviour. Over the course of infection, the food intake of infected male mice decreased from the onset of infection until 11 days post-infection (dpi), after which it increased to that of naïve males, before dropping again at 25 dpi (**Figure 1E**). Infected female mice displayed a brief, but non-significant, reduction in food intake at 7 dpi, but otherwise maintained a similar feeding behaviour profile to their wild type counterparts (**Figure 1F**). Taken together, these data suggests that weight loss during experimental trypanosomiasis occurs in a sex-dependent manner and may be associated, in males, to changes in feeding behaviour.

We next explored the impact of *T. brucei* infection on the adipose tissue. In addition to changes in feeding behaviour^23^, weight loss is typically associated with reductions in adipose tissue mass^24^. We were particularly interested in the iWAT, which is analogous to the scWAT tissue in humans, as this constitutes an important parasite niche for disease transmission, especially in asymptomatic carriers^16^. Previous reports have shown that colonisation of the gonadal white adipose tissue (gWAT) is associated with weight loss and reduction in adipose mass^6^, but the effect on iWAT, which is anatomically and functionally distinct from gWAT (**Figure 2A**), was not investigated. We first determined the presence of parasites in the iWAT and gWAT by histological analysis (**Figure 2B**) and, as expected, detected trypanosomes in both tissues. Next, we quantified trypanosome genomic DNA in the iWAT and gWAT as a proxy for determining parasite density. We found that in male mice there were fewer parasites in the iWAT compared with the gWAT, but that there were no differences between these depots in females (**Figure 2C**). Our data are consistent with previous reports^6^, and highlights that the iWAT also harbours a population of parasites that, given their proximity to the skin, are important for forward parasite transmission^16^.

**Figure 2.**
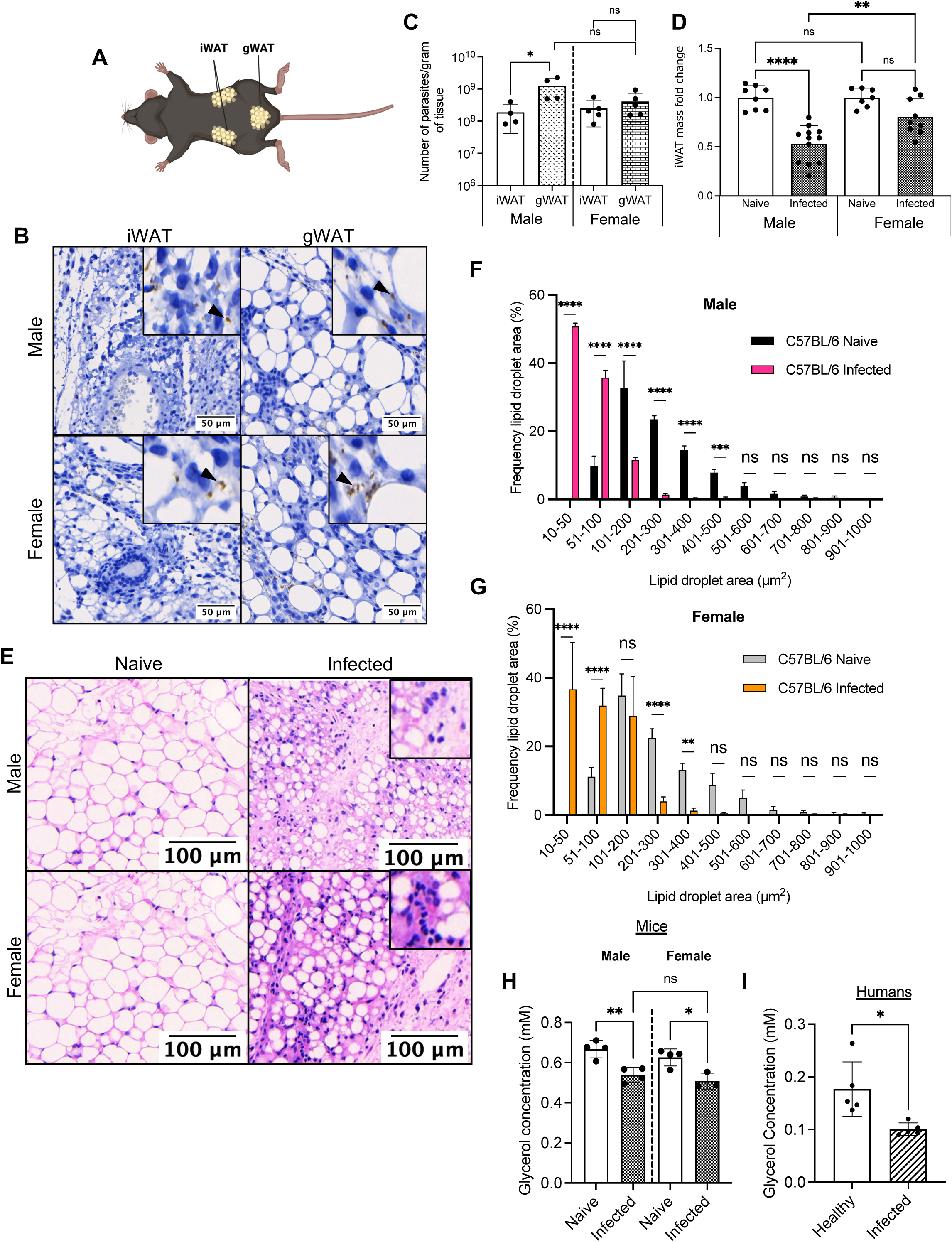
*Trypanosoma brucei* infection leads to reductions in iWAT mass and lipid content, as well as impairment of adipose tissue function. **(A)** Schematic highlighting the anatomical location of the inguinal white adipose tissue (iWAT) and gonadal white adipose tissue (gWAT). **(B)** Histological analysis of the iWAT and gWAT trypanosome colonisation, using HSP70 staining. **(C)** Parasite burden of iWAT and gWAT. Parasite density in the iWAT, which was measured by RT-qPCR of genomic DNA. A comparison was made with gonadal white adipose tissue (gWAT), to understand if the iWAT is also highly colonised. **(D)** iWAT mass at 25 days post-infection or in naïve mice. iWAT was dissected and weighed before normalising to body weight, to account for variation between biological replicates. Symbols indicate the number of biological replicates collected from two independent experiments **(E)** Representative histological H&E staining of iWAT showing adipocyte lipid droplets and immune infiltrate. Insets highlight likely immune cell infiltrate. **(F)** Analysis of lipid droplet area (µm^2^) in naïve and infected males and **(G)** females. *N*=6 biological replicates per group, from two independent experiments. Lipid droplets were measured from 3 distinct areas in each image and then combined for each biological replicate. **(H)** Circulating glycerol concentration in naïve vs. infected male and female mice. *n*=4 biological replicates per group. **(I)** Circulating glycerol concentration in healthy vs. infected patients. *N*=5 patients per group and each group is a mix of male and female. For **C**, **D, H** and **I**, data were analysed using a two-tailed Student t test. For **F** and **G**, data were analysed using a two-way ANOVA with Sidak post-hoc testing. Data for all panels are expressed as mean ±SD. \**p*<0.05, \*\**p*<0.01, \*\*\**p*<0.001, \*\*\*\**p*<0.0001, ns = non-significant.

Following our observations of weight loss and the presence of trypanosomes in the iWAT, we then proceeded to characterise the impact of infection on this adipose tissue depot. When normalised to bodyweight, we found that infection led to a significant reduction in the iWAT mass of male mice (**Figure 2D**). Female mice also experienced a reduction in iWAT mass, but this was not significant. This raised the question of whether the reduction in mass is due to a loss of lipid content and reduction in adipocyte size (hypotrophy). To address this, we performed Haematoxylin and Eosin (H&E) staining of iWAT at 25 dpi. We found that the iWAT of males and female mice infected with *T. brucei* undergoes hypotrophy, with concurrent infiltration of immune cells (**Figure 2E**). Furthermore, we found a differential effect on the morphometric properties of the iWAT adipocytes in response to *T. brucei* infection. In infected male mice, there was a significant decrease in lipid droplet size, with the majority of lipid droplets ranging from 10-200 µm^2^, compared with naïve controls where lipid droplet size was larger, with the majority of droplets ranging from 51-900 µm^2^ (**Figure 2F**). In infected female mice, there was a less dramatic shift in lipid droplet size, with the majority of lipid droplets ranging from 10-400 µm^2^, compared with naïve controls where lipid droplet sizes ranged from 51-1,000 µm^2^ (**Figure 2G**). The morphometric analyses of iWAT in response to infection are indicative of tissue wasting. Thus, we questioned whether *T. brucei* infection impacted adipose tissue function, using systemic glycerol levels as a proxy for adipose tissue function^25^. In both infected male and female mice, the circulating glycerol levels were significantly reduced compared to naïve controls (**Figure 2H**), consistent with a global impact of infection on adipose tissue function. The changes in glycerol levels observed in experimental infections were also replicated in the serum of stage II HAT patients from the towns of Boffa, Forécariah and Dubréka in Guinea, suggesting that the adipose tissue dysfunction induced by infection also occurs in humans (**Figure 2I**). Taken together, these findings highlight that *T. brucei* infection is associated with significant iWAT wasting and that this, in turn, is associated with impaired tissue function in both mice and humans.

### *T. brucei* iWAT colonisation results in a hypometabolic state in the murine iWAT

To better understand the iWAT response to infection, and to identify potential drivers of tissue wasting, we performed bulk transcriptomic analysis of iWAT harvested at 25 dpi from both sexes and included naïve controls for comparison. Principal component analysis (PCA) revealed high levels of variance between naïve and infected males, but less variance between naïve and infected females (**Figure 3A**), potentially indicating a higher transcriptional response in the iWAT of male mice compared to female mice. Indeed, compared with naïve controls, differential expression analysis revealed upregulation of 3,828 and 3,177 genes with a log_2_Fold change >0.5 and an adjusted *p* value (*padj*) of <0.01 in infected male and female mice, respectively (**Figure 3B; Supplementary Table 1**). In contrast, compared with naïve controls, this analysis revealed downregulation of 3,332 and 2,606 genes a log_2_Fold change >0.5 and an *padj* of <0.01 in infected male and female mice, respectively (**Figure 3B; Supplementary Table 1**). To explore this dataset further, we performed pathway enrichment analyses, enriching for Kyoto Encyclopaedia of genes and genomes (KEGG) terms. We first observed that in both males and females, the majority of upregulated pathways were associated with immune and inflammatory processes (**Figure 3C and D**). Many genes within these pathways related to MHC class II antigen presentation (e.g. *H2-DMa*, *H2-DMb1*, *H2-Ob*), inflammatory cytokines (*e.g. Tnf*, *Ifng*, *Il1b*), and complement activation (e.g. *C2*, *C3*, *C4b*) (**Supplementary Table 2**). When we explored downregulated pathways, we found that the majority of these were related to metabolism (**Figure 3E and F**), including amino acid degradation (e.g. *Abat*, *Acat1*, *Acat2*) and propanoate metabolism (e.g. *Sucla2*, *Acox1*, *Acads*) (**Supplementary Table 2**). Unexpectedly, we also observed that *T. brucei* infection led to a downregulation of the lipolysis pathway in males and females (**Supplementary Figure S2**), with canonical genes such as *Pnpla2*, *Fabp4,* and *Lipe* being significantly downregulated (**Supplementary Figure S2)**. Since lipolytic genes were downregulated, we questioned whether the iWAT was becoming more thermogenic, which could, in turn, lead to a decrease in mass. However, when exploring our data, we also found that canonical genes associated with thermogenesis, such as *Ucp1*^26^, were downregulated in infected males and females (**Supplementary Table 2**), Moreover, expression of genes associated with UCP1-independent thermogenesis, including *Atp2a1*, *Atp2a2*, and *Atp2a3*^27^ were not altered during infection (**Supplementary Table 2**), suggesting that thermogenesis is not activated in this context. These results, together with an overall decrease in circulating glycerol levels in serum, indicate that chronic *T. brucei* infection results in a hypometabolic state in mice (and likely in humans) to conserve energy^28^.

**Figure 3.**
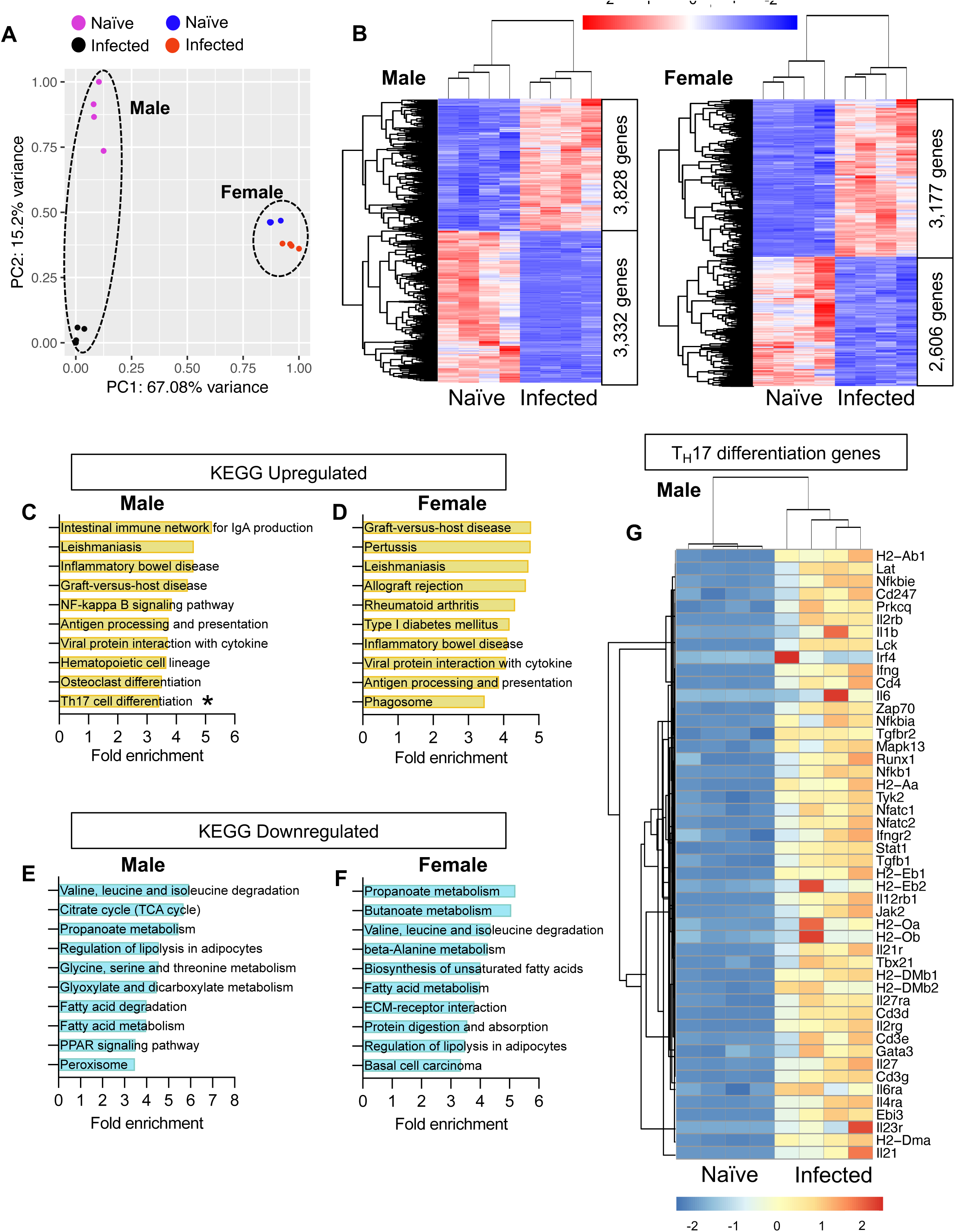
Transcriptomic analyses indicate a stronger T_H_17 response in the iWAT of males than females. **(A)** Principal component analysis of bulk transcriptomic data from male and female naïve and infected mice. **(B)** Heatmaps, clustered by Euclidean distance, of differentially expressed genes in the iWAT of naïve vs infected male and female mice. (**C and D**) Pathway enrichment analysis of upregulated and downregulated genes in male mice. (**E and F**) Pathway enrichment analysis of upregulated and downregulated genes in female mice. **(G)** Heatmap of T_H_17 transcripts in male mice. *n*=4 biological replicates per group.

### *T. brucei* iWAT colonisation induces an upregulation of genes associated with T_H_17 T cell differentiation

Our transcriptomic analyses also revealed specific enrichment of T cell-related transcripts in male but not female mice. For example, we found *Cd3d*, *Cd3e*, and *Cd247* were significantly upregulated in the iWAT of infected male mice but exhibited no changes in infected females (**Supplementary Table 2**). T_H_1- and T_H_2-related transcripts included *Nfatc1*, *Cd4*, *Runx3,* and *Gata3*, suggesting that T_H_1 cells may be a significant contributor to interferon gamma (IFNγ) production during infection in the adipose tissue. Additionally, we also detected a significant upregulation of genes associated with differentiation of T_H_17 effector cells, including *Irf4*, *Cd4*, *Il21r*, and *Il6ra* (**Figure 3G**).

Based on our observations, we hypothesised that T_H_17 cells are important for the adipose immune response to infection. To quantify the different populations of CD4^+^ T cells, including T_H_17 cells, present in the iWAT during chronic *T. brucei* infection, we utilised mass cytometry by time of flight (CyTOF), enabling us to gain as much information as possible from the wasted adipose tissue. Consistent with previous studies^8^, we also found an expansion of macrophages, granulocytes, and dendritic cells in the iWAT of infected males and female mice (**Figure 4A-4D**). The B cell subset was decreased specifically in male mice during infection compared to naïve controls, without noticeable changes in female mice (**Figure 4E**). Regarding the T cell effector population, we observed an increase in the proportion of CD3ε^+^ TCRβ^+^ CD4^+^ T cells in the iWAT of infected mice (**Figure 4F**) and, in particular, we identified an expansion of CD44^+^ CD69^+^ CD4^+^ T effector (Teff) cells (**Figure 4G**). Whilst the frequency of CD4^+^ T cells increased in both males and females, the iWAT of infected females contained a higher proportion of Teff cells compared with males. The expanded Teff cells displayed an elevated production of interferon gamma (IFNγ; **Figure 4H**), suggesting that some of these cells are polarised towards a T_H_1 phenotype. Furthermore, there was a significant expansion of IL-17A-producing Teff cells (**Figure 4I**), consistent with our transcriptomics dataset. Interestingly, in addition to TNFα and IFNγ, IL-17A was also elevated in the serum of infected mice and compared to naïve controls (**Figure 4J**), and in second stage HAT patients compared to healthy controls (**Figure 4K**), demonstrating that IL-17A elevation is conserved mice and humans infected with *T. brucei*. Taken together, our data demonstrate a significant expansion of IL-17-producing T cells in the iWAT in response to *T. brucei* infection, and may be associated with the induction of a hypometabolic state.

**Figure 4.**
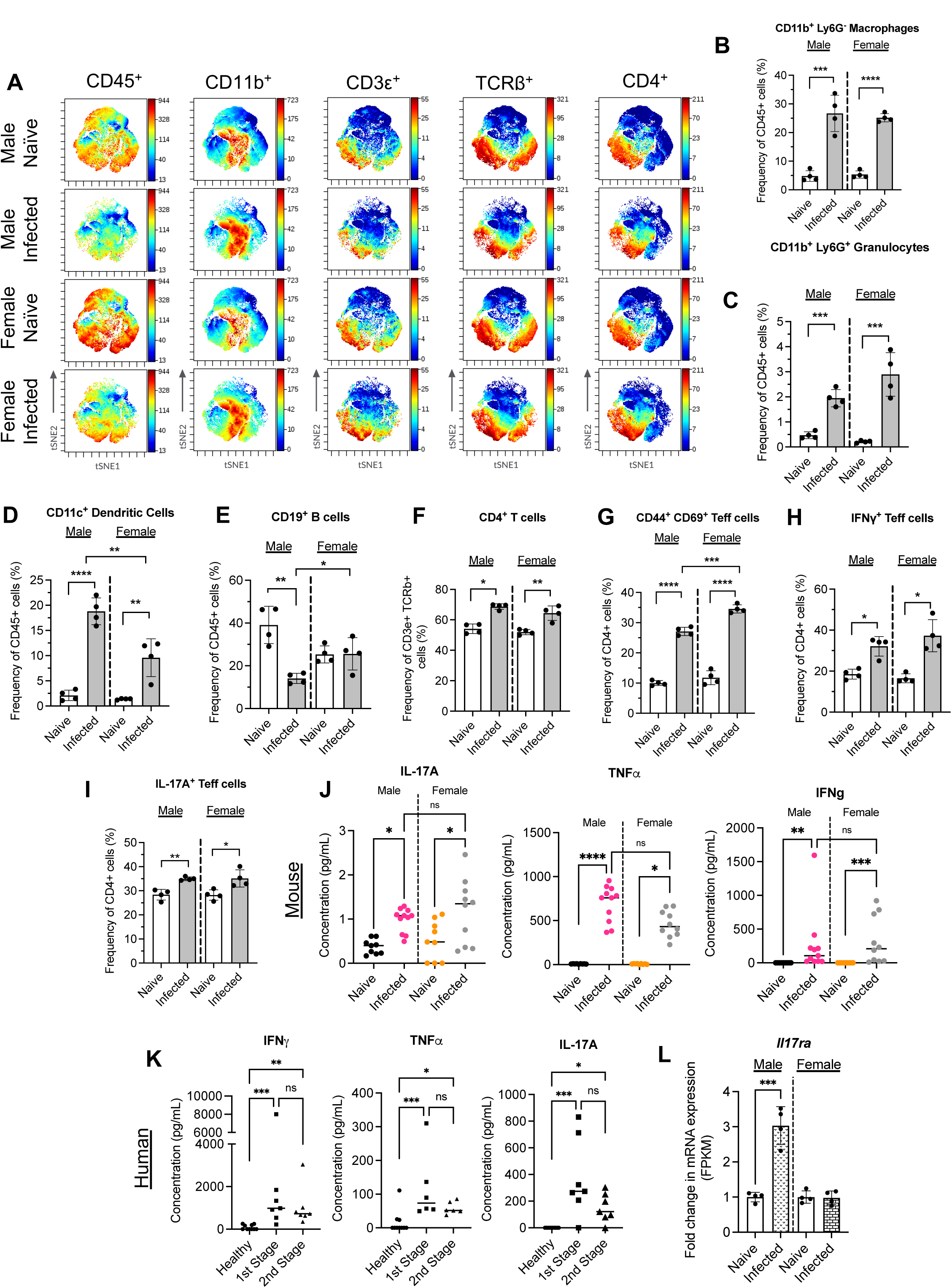
Elevated circulating IL-17A is associated with increased IL-17 receptor A (IL-17RA) in male mice during *T. brucei* infection. **(A)** t-distributed stochastic neighbour embedding (t-SNE) plots of broad macrophage and T cell populations. We gated on CD45^+^, CD3ε^+^, TCRβ^+^, CD4^+^ cells. (**B**) Measurements of the proportions of macrophages, **(C)** granulocytes, **(D)** dendritic cells **(E)** and B cells. **(F)** Measurements of the proportions of CD4^+^, **(G)** T effector (Teff), **(H)** IFNγ-producing and **(I)** IL-17A-producing T cells. Measurements of circulating serum IL-17A, TNFa, and IFNγ in mice **(J)** and IFNγ, TNFa and total IL-17 in humans **(K)**. **(L)** Expression of interleukin-17A receptor (*Il17ra*) mRNA in infected male and female mice. Mouse cytokine data were collected from samples taken across three independent experiments. Data were tested for normal distribution and analysed by either one-way ANOVA or a Kruskal Wallis test. Biological replicates are indicated by symbols for each panel. Data for all panels are expressed as mean ±SD. \**p*<0.05, \*\**p*<0.01, \*\*\**p*<0.001, \*\*\*\**p*<0.0001, ns = non-significant.

### IL-17A/F is critical for controlling bodyweight and pathology during *T. brucei* infection

Our results so far indicate that chronic iWAT infection leads to an expansion of local T_H_17 effector T cells in male mice. However, the role of IL-17 in the control of *T. brucei* infection, or in the local infection-induced pathology in the iWAT, has not been explored to date. To test this, we used a global *Il17a/f* knockout mouse, which is deficient in both IL-17A and IL-17F^29^ (**Figure 5A** **and Supplementary Figure S3A**). Systemically, we observed that the first peak of parasitaemia was similar between wildtype and *Il17af^-/-^*male mice, with a non-significant increase in parasitaemia post-day 13 onwards in the *Il17af^-/-^*mice (**Figure 5A**), which may suggest that IL-17A/F plays a role in controlling systemic parasite numbers during chronic infection. In contrast, the first peak of parasitaemia in *Il17af^-/-^* females was lower than in C57BL/6 mice, and the second peak of parasitaemia was delayed (**Supplementary Figure S3A)**. Deficiency of *Il17a/f* was also associated with an earlier onset and increased severity of clinical symptoms in both males (**Figure 5B**) and females (**Supplementary Figure S3B**). *Il17af^-/-^ m*ale and female mice started to exhibit clinical symptoms (piloerection and hunching) from 3 and 7 dpi, respectively, whereas wild type mice started to experience these symptoms between 12 and 15 dpi. Unlike bloodstream parasite numbers, the parasite burden of the major adipose tissue depots does not appear to be influenced by IL-17A/F in either sex (**Figure 5C** a**nd Supplementary Figure S3C and S3D**), indicating that IL-17A/F is dispensable for controlling systemic of local parasites burden in the adipose tissue.

**Figure 5.**
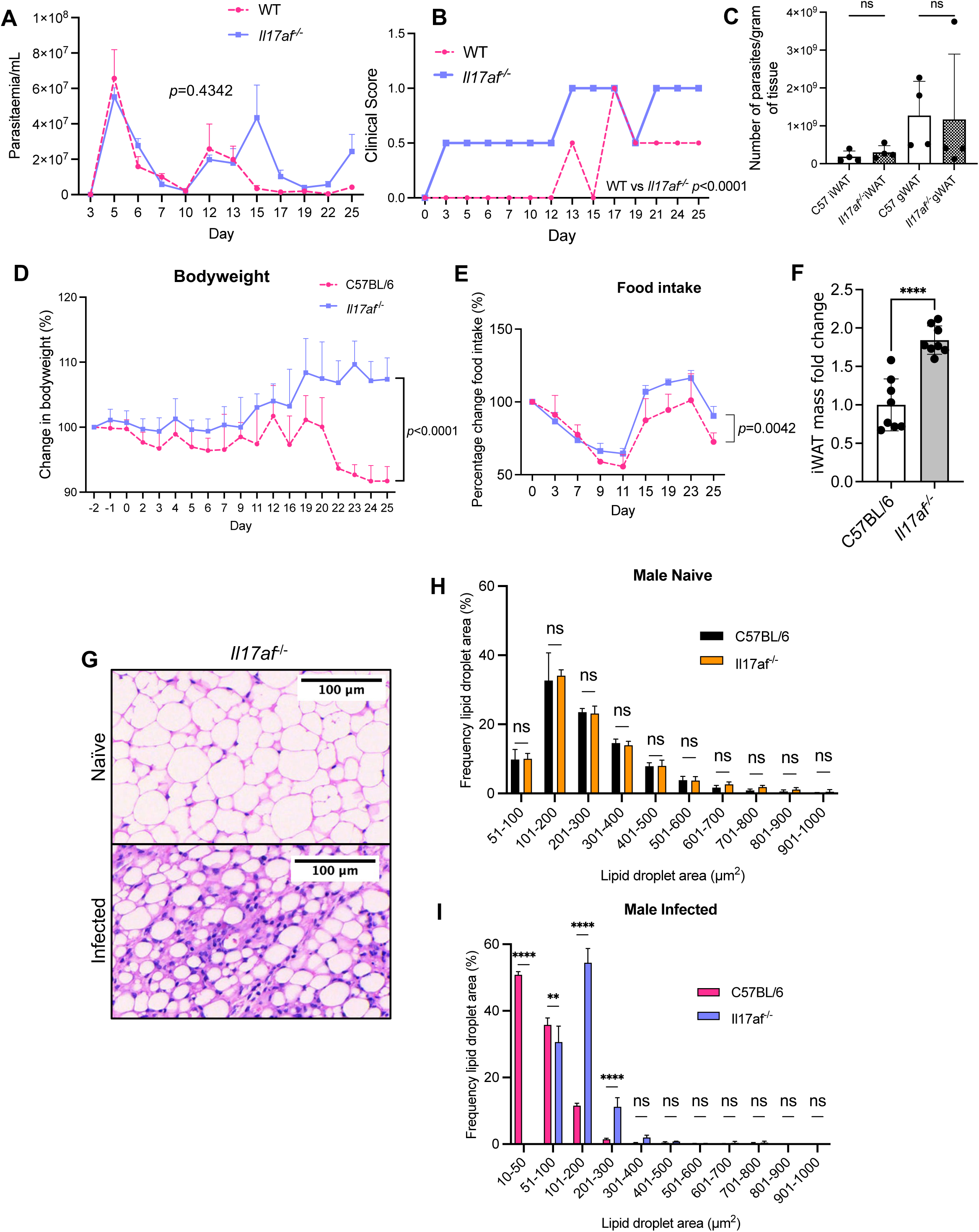
IL-17A/F limits clinical symptoms during *T. brucei* infection and drives weight adipose tissue wasting in male mice. (**A**) Number of parasites per mL of blood in *Il17af^-/-^* vs wild type male mice, which was measured using phase microscopy and the rapid “matching” method^53^. (**B**) Comparison of clinical scores between C57BL/6 and *Il17af^-/-^* mice. **(C)** Parasite burden of inguinal WAT (iWAT) and gonadal WAT (gWAT). **(D)** Percentage changes in bodyweight of wild type and *Il17af^-/-^* male mice over the course of infection. *n*=7 mice per group across two independent experiments. **(E**) Changes in gross food intake over the course of infection. Each data point represents 2 cages (*n*=3-4 mice per condition) from 2 independent experiments. **(F)** iWAT mass in infected C57BL/6 and *Il17af-/-* male mice. **(G)** H&E staining of iWAT from male *Il17af^-/-^* mice. (**H and I**) Analysis of lipid droplet area (µm^2^) in naïve and infected male mice. Lipid droplets were measured from 3 distinct areas in each image and then combined for each biological replicate. Data for C57BL/6 mice in panels **A, B, D** are taken from Figure 1. Data for C57BL/6 mice in panels **H** are taken from Figure 2. Data were analysed using two-way repeated measures ANOVA with Sidak post-hoc testing or a one-way ANOVA with Tukey post-hoc testing. Data points represent biological replicates. With the exception of panels **C** and **G**, data are from 2-3 independent experiments. Data are expressed as ±SD. \*\**p*<0.01, \*\*\*\**p*<0.0001.

Having established that IL-17A/F signalling is critical for controlling pathology during *T. brucei* infection, we next examined the role IL-17A/F in controlling local iWAT pathology. For this, we monitored the weight of both male and female mice during the course of infection. Unlike C57BL/6 male mice, infected *Il17af^-/-^*males increased their bodyweight (∼8-10%) during infection (**Figure 5D**). Infected female *Il17af^-/-^* mice also gained weight during infection (∼8-19%) in a similar pattern to C57BL/6 females, although similarly to males, IL-17A/F-deficient females gained more weight than their wild type counterparts (**Supplementary Figure S3E**). Previous reports have shown that when administered to wild type naïve mice under homeostatic conditions, IL-17A suppresses food intake^30^. Therefore, given the potential link between IL-17 signalling, bodyweight, and food intake, we next measured gross food intake in infected C57BL/6 and *Il17af^-/-^* mice. We observed that up until 9 dpi, wild type and *Il17af^-/-^* male mice reduced their food intake at the same rate (**Figure 5E**). However, after 11dpi, *Il17a^-/-^* mice started to increase their food intake above that of the wild types. Indeed, food intake for infected *Il17af^-/-^*mice was higher between 15 and 23 dpi compared with at the onset of infection. Unlike male mice, food intake was indistinguishable between infected female *Il17af^-/-^* and wild type mice (**Supplementary Figure 3SF**). Together, our results suggest that IL-17A/F potentially regulates bodyweight by altering food intake in male mice during infection, but not female mice. Importantly, the weight gain in females was also associated with increased splenomegaly in *Il17af^-/-^* mice, compared with their wild type counterparts or to male mice, thus indicating that splenomegaly by itself does not account for the increase in bodyweight observed in male mice (**Supplementary Figure S3G and S3H**). Related to the differences in bodyweight, we also found that upon infection, *Il17af^-/-^* knockout male mice also retained more of their iWAT mass compared with their wild type counterparts (**Figure 5F**), suggesting that they experienced less wasting. In contrast, although naïve *l17af^-/^* female mice have a higher mass of iWAT than their wild type counterparts, they experience similar levels of wasting as wild type females upon infection (**Supplementary Figure S3I**), supporting our hypothesis that the effects of IL-17A/F are sex-dependent.

To understand whether global deficiency of *Il17af^-/-^* impacts iWAT lipid content, we performed H&E staining on the iWAT and measured lipid droplet size in naïve and infected male (**Figure 5G** **-5I**) and female (**Supplementary Figure S3J to S3L**) mice and compared these with wild type animals. The iWAT adipocyte size was indistinguishable between genotypes in naïve animals, indicating that IL-17A/F does not affect lipid droplet size under homeostasis. However, upon infection, *Il17af^-/-^* males retained a higher frequency of larger lipid droplets than wild type males (**Figure 5I**). When comparing this to female mice, we found that deletion of *Il17af^-/-^* had no impact on the range of lipid droplet sizes during *T. brucei* infection (**Supplementary Figure S3L**). Our results demonstrate that IL-17A/F signalling drives iWAT wasting and lipid usage in adipocytes in males, but not in females, during *T. brucei* infection.

### Single cell analysis of the iWAT stromal vascular fraction reveals a distinct population of IL-17-producing CD27^-^ Vγ6^+^ T cells in the iWAT of infected mice

To further understand the events leading to iWAT wasting in response to *T. brucei* infection, we conducted single cell RNA on the iWAT, now focusing on male mice since this is where we observed both upregulation of the IL-17A receptor (**Figure 4L**) and changes in bodyweight (**Figure 5D**). As we wanted to understand the early processes leading up to the loss of iWAT tissue mass, we infected mice for 7 days before harvesting the iWAT. Following iWAT dissociation and scRNAseq analysis, we obtained a total of 46,546 high-quality cells with an average of 1,296 genes and 20,209 reads per cell (**Supplementary Table 3**). These cells were broadly classified into 18 clusters (**Figure 6A**) based on common markers associated with these clusters. The predominant cell type in this dataset was immune cells, with eight B cell clusters, four T cell clusters, an NK cell cluster, two macrophage clusters, and a plasmacytoid dendritic cell (pDC) cluster (**Figure 6A**). In addition to the cells within the immune compartment, we identified two stromal cell clusters comprised of preadipocytes. Of the cell types we identified, we observed that several expanded in the iWAT of infected mice, including Tregs, T cell 1, Transitional B cell 4, NK cells, and Macrophage 2, whereas other cell populations remained similar in numbers between conditions, or contracted in the case of preadipocytes T cell 2, several B cell clusters, and pDCs (**Figure 6B**). Of the B cells, we classified four clusters as transitional B cells, based on high expression of *Cd79a*, *Cd79b*, and *Ighd*, and we classified plasma cells based on these markers with the addition of *Jchain, Ighm,* and *Sdc1* (**Figure 6C**). We also identified a cluster of germinal centre (GC)-like B cells, which were classified based on expression of *Aicda* and *Pcna*. T cells were classified as CD8^+^ T cells (*Cd3d^+^*, *Cd3e^+^*, *Cd8a^+^*, *Trac^+^*), Tregs (*Cd3d^+^*, *Cd3e^+^*, *Cd4^+^*, *Foxp3^+^*), NK cells (*Cd3d^+^*, *Cd3e^+^, Gzma^+^, Gzmb^+^, Nkg7^+^*), or T cell 1 and T cell 2 (*Cd3d^+^*, *Cd3e^+^*, *Trac^+^*). The myeloid compartment was composed of two macrophage subclusters, classified as Macrophage 1 (*Lyz2^+^*, *Fcer1g^+^*, *S100a4^+^*) and Macrophage 2 (*Ccl5^+^, S100a4^+^, Tmem176a^+^, Tmem176b^+^*). Finally, we classified pDCs based on high expression of *Fcer1g*, *Lgals1*, *Siglech*, and *Runx2*. Given the prominent role of genes associated with T_H_17 differentiation detected in the iWAT from infected mice detected by bulk transcriptomics (Figure 3), we next proceeded to resolve the intrinsic heterogeneity of the T cell subset associated with the iWAT. For this, we subset and reanalysed the T cell compartment separately (**Figure 6D**). This resulted in five T cell subclusters, which consisted mainly of Tregs (*Cd4, Icos, Foxp3*), CD8^+^ T cells (*Cd8a*, *Cd8b1*), activated NK cells (*Nkg7, Klr1d1, Gzma, Gzmb, Nr4a1^+^*), a cluster of activated and replicative T cells with evidence of TCR engagement (*Top2a^+^, Mki67^+^, Hist1h1b^+^, Nr4a1^+^*), and IL-17A^+^ Vγ6^+^ cells (*Tcrg-C1^+^, Rorc^+^, Cd163l1^+^, Il17a^+^*) (**Figure 6E**). To validate the *in silico* prediction that there is an expansion of IL-17A^+^ cells in the murine iWAT in response to infection, we next measured IL-17A expression using a murine reporter line, in which IL-17A expression is coupled to GFP expression (IL-17A^GFP^). We observed that at 7 days post-infection, there was a significant expansion of IL-17A^+^ cells (**Figure 6F**), including T_H_17 (CD45^+^, CD3ε^+^, CD4^+^, GFP^+^) cells (**Figure 6G**). Although we did not observe a change in the frequency of γδ T cells (**Figure 6H**), consistent with our *in silico* predictions, we observed a significant increase in the frequency of IL-17A-producing CD27^-^ γδ T cells (**Figure 6I**) and a concomitant decrease in the frequency of IFNγ-producing CD27^+^ γδ T cells (**Figure 6J**). Together, these findings demonstrate a dominant IL-17-driven response in the murine iWAT in response to *T. brucei* infection. Our data also demonstrate that IL-17A is derived from multiple sources including local T_H_17 cells and CD27^-^ Vγ6^+^ T cells.

**Figure 6.**
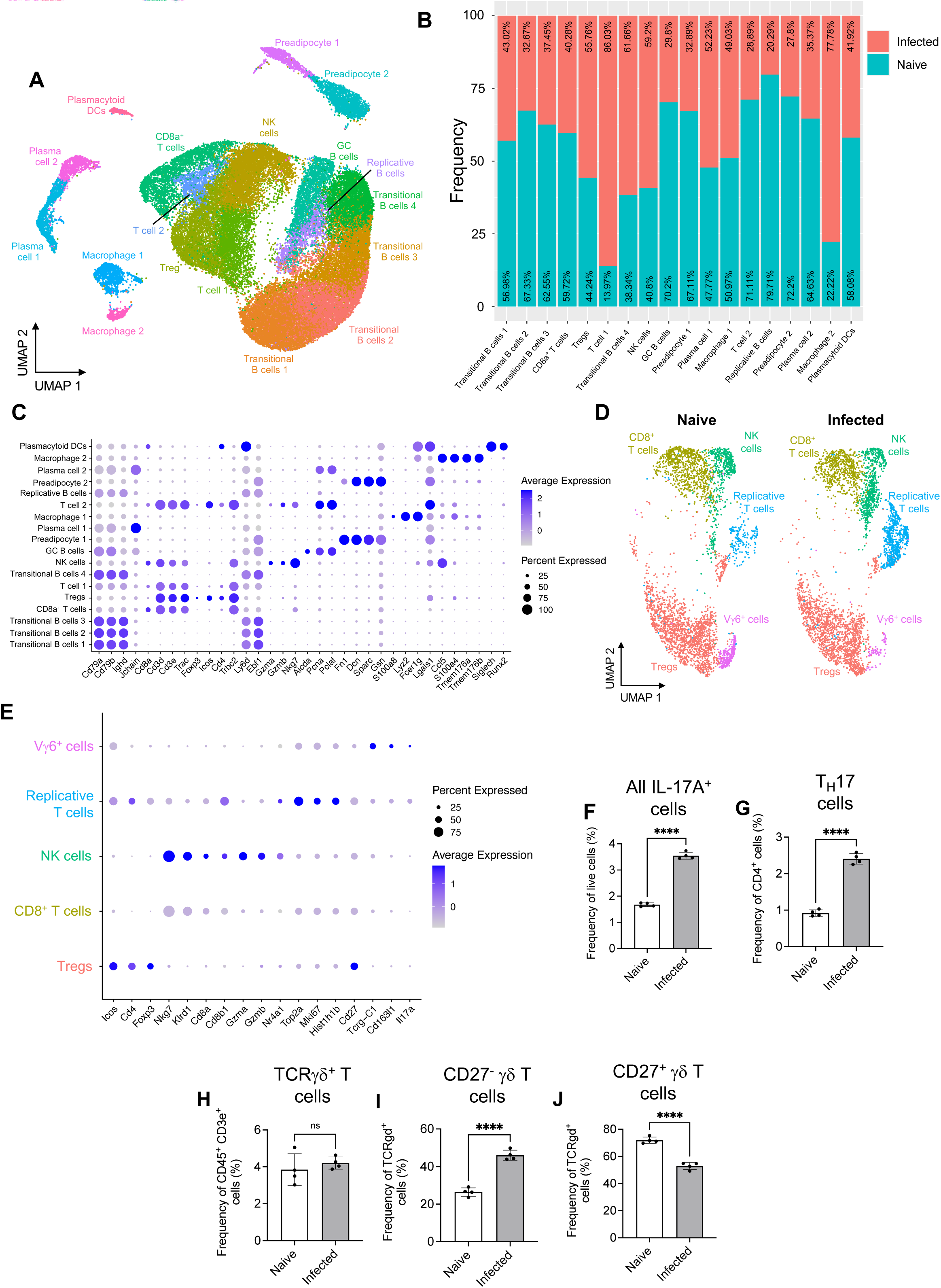
*T. brucei* infection leads to expansion of IL-17A^+^ cells in the iWAT during early infection. (**A**) Uniform manifold approximation and projection (UMAP) of 46,546 high-quality cells from single cell RNA sequencing (scRNAseq) dataset. (**B**) Frequency plot showing changes in cell frequency under naïve or infectious conditions. (**C**) Dot plot representing the expression levels of top marker genes used to catalogue the diversity of cells within the dataset. The side of the dots represent the percentage of cells that express a given marker, and the colour intensity represent the level of expression. (**D**) UMAP of subset T cell clusters. (**E**) Dot plot representing the expression levels of top marker genes used to categorise T cell populations. (**F**) Flow cytometry analysis of the proportion of all live IL-17A^+^ cells. Frequency of (**G**) T_H_17, and (**H**) TCRγδ^+^ T cells in the iWAT. Frequency of (**I**) CD27^-^ and (**J**) CD27^+^ γδ T cells in the iWAT. Single cell data are comprised of one technical replicate per condition, each containing pooled cells from the iWAT of 5 male mice replicate. For flow cytometry data, data, biological replicates are indicated by symbols for each panel, and data were analysed using a student’s t-test. Data are expressed as ±SD. \**p*<0.05, \*\*\*\**p*<0.0001, ns=non-significant.

### *T. brucei* infection results in broad transcriptional changes in iWAT preadipocytes, including upregulation of the IL-17A receptor and lipolysis

In addition to detecting a range of IL-17A-producing cells by scRNAseq, we also wanted to understand which cells may be responding to this cytokine during *T. brucei* infection. To this end, we measured IL-17A receptor (*Il17ra*) expression across all clusters and found that it was exclusively upregulated in preadipocytes during infection (**Figure 7A**), indicating that of all the cells within the iWAT stroma, preadipocytes are likely to be responsive to IL-17A. However, we were unable to resolve the preadipocyte heterogeneity within the iWAT, or their responses to infection, so we reclustered these cells and reanalysed them. Using previous reports to annotate the preadipocytes^31, 32^, and after reclustering the population of iWAT preadipocytes (**Figure 6A**), we identified five distinct populations (**Figure 7B**), four of which expressed canonical mesenchymal markers including *Ly6a, Pdgfra*, and *Cd34* (**Figure 7C**)^32^. These mesenchymal subclusters encompassed interstitial preadipocytes (*Dpp4, Pi16, Bmp7*), located in the interstitium, which are known to be poised to migrate into the iWAT to differentiate into mature adipocytes when needed^32^, a committed preadipocyte cluster (*Col4a1, Col4a2, Col15a1, Fabp4, Plin2*), which represent a transitional state between stem-like preadipocytes and mature adipocytes, and an adipogenesis regulatory cell cluster (*Fmo2, F3, Clec11a*), which are able to suppress adipogenesis *in vivo*^33^. Lastly. We also found a cluster of mature adipocytes (*Cd36, Fabp4, Pparg*) (**Figure 7B and 7C**). Importantly, the mature adipocyte populations were exclusively detected in the naïve samples and not in infected controls, perhaps due to the wasting associated with infection. During infection, we noted an increase in the frequency of the interstitial preadipocyte 1 cluster compared to naïve controls (**Figure 7D**), resulting in a decrease in the frequency of other subsets, including mature adipocytes, which might reflect the histological findings associated with iWAT wasting. Of the preadipocyte populations that we identified, we noted that *Il17ra* expression was robustly expressed in both interstitial preadipocyte clusters, and to a lesser extent in the committed preadipocyte cluster, (**Figure 7E**), but not in adipogenesis-regulatory cells or mature adipocytes, suggesting that both interstitial and committed preadipocytes are able to sense IL-17A/F signalling in response to infection. Using *in silico* gene module scoring to assess expression of genes involved in lipolysis (e.g. *Pnpla2, Plaat3, Mgll, Fabp4*), we identified that responses to chronic *T. brucei* infection were observed in committed preadipocytes and adipogenesis-regulatory cells (**Figure 7F**). This possible elevation of lipolysis is in contrast to our data from later timepoints of infection, where we found evidence of decreased lipolysis (**Figure 2H, 2I, 3G, and 3H**). Glycerol, which is released through lipolysis can also enter the glycolysis pathway, where it can contribute to either energy generation through entry into the tricarboxylic acid (TCA) cycle, or it can enter the pentose phosphate pathway (PPP) and can support inflammation^34^. We therefore looked at the expression of genes associated with glycolysis and the TCA cycle (**Supplementary Figure S4A**). We found that genes associated with glycolysis and the downstream generation of lactate (*Hk2, Ldha*) were upregulated during infection, across all preadipocyte populations. We also found upregulation of key TCA cycle genes, including those encoding for succinate dehydrogenase (*Sdhb*) and malate dehydrogenase (*Mdh1, Mdh2*) during infection, which could contribute to reactive oxygen species (ROS) generation. Finally, we explored genes in the pentose phosphate pathway (**Supplementary Figure S4B**) and found upregulation of key genes within this pathway (including *Aldoa, Gpi1*). Taken together, these results demonstrate that during early infection time points, interstitial and committed preadipocytes upregulate gene pathways associated with IL-17 signalling and lipolysis.

**Figure 7.**
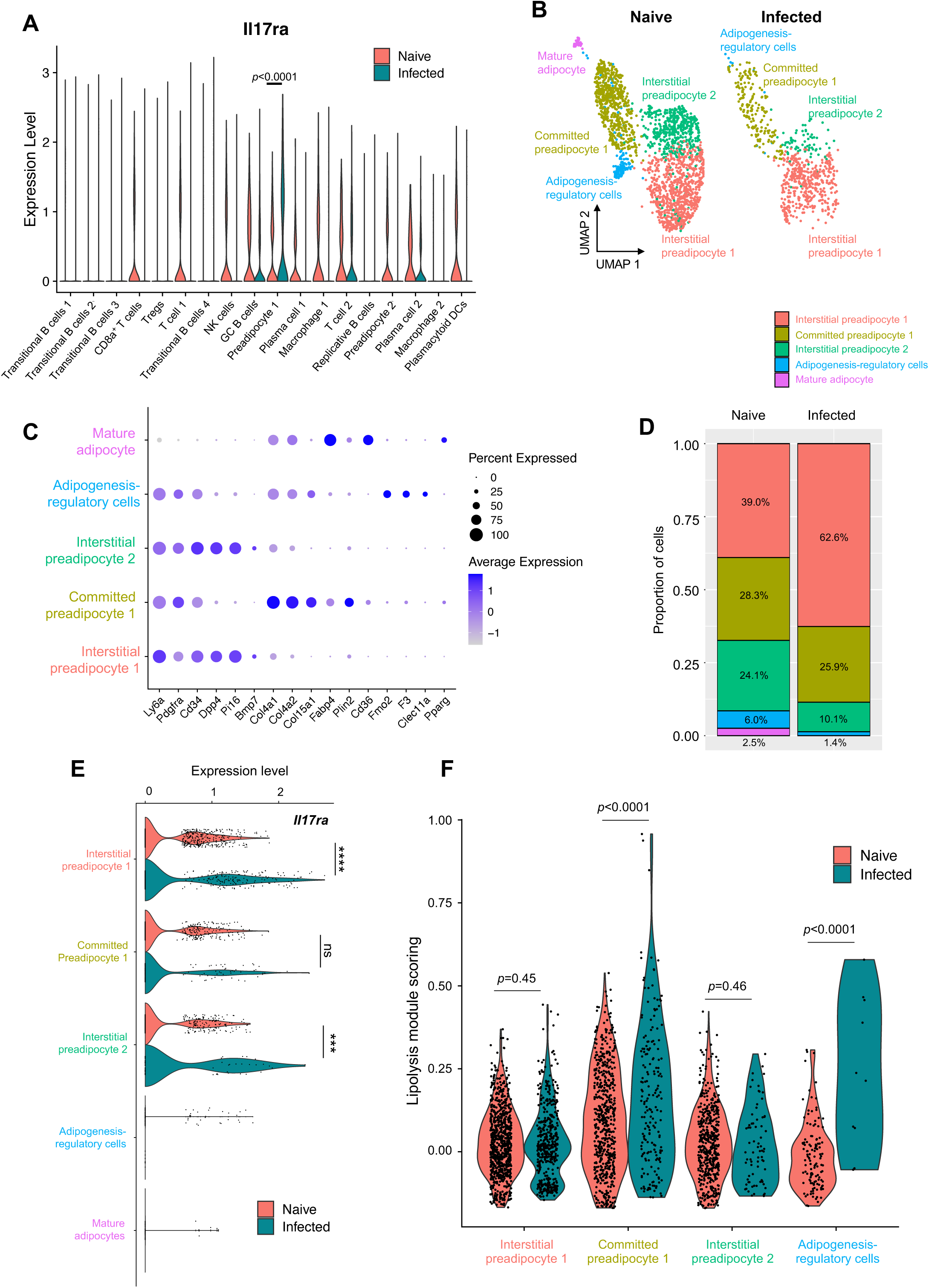
*T. brucei* infection leads to upregulation of the IL-17A receptor and genes associated with lipolysis in preadipocytes. (**A**) Violin plot showing the expression level of the IL-17A receptor (*Il17ra*) across all cell populations in our scRNAseq dataset. (**B**) UMAP of subclustered preadipocytes. (**C**) Dot plot showing expression of genes used to identify specific populations of preadipocytes. (**D**) Frequency plot of preadipocyte populations. (**E**) Violin plot showing expression of *Il17ra* in specific preadipocyte populations. (**F**) Lipolysis gene module score of genes typically associated with lipolysis across all preadipocyte subsets detected in (**B**). Single cell data are comprised of one technical replicate per condition, each containing pooled cells from the iWAT of 5 male mice replicate.

### IL-17A/F signalling on adipocyte is critical for controlling tissue wasting and local parasite burden in the iWAT

Our scRNAseq dataset clearly demonstrates that both interstitial and committed preadipocytes upregulate *Il17ra* expression in response to *T. brucei* infection, rendering them able to sense IL-17A/F locally. Since committed preadipocytes are on a fixed trajectory to a mature adipocyte lineage^35^, we asked whether IL-17A/F signalling through adipocytes plays a role in driving changes in bodyweight, iWAT mass and/or iWAT lipid content. To address this, we generated mice with a constitutive deficiency of *Il17ra* in white adipocytes, across all white adipose tissue depots (*Adipoq^Cre^*x *Il17ra^fl/fl^*) and infected them alongside C57BL/6 controls, housing them in single-housing. We observed that adipocytes from *Adipoq^Cre^*x *Il17ra^fl/fl^* mice maintained their bodyweight (and at points increased their weight) throughout the course of infection compared to wildtype controls, which lose bodyweight, in particular after 20dpi (**Figure 8A**). These results mirrored the results obtained from the *Il17af^-/-^*, which were also protected from the infection-induced weight loss (**Figure 7D** **and Supplementary Figure S5A**). Additionally, we observed that both *Il17af^-/-^*and *Adipoq^Cre^* x *Il17ra^fl/fl^* mice had a higher food intake over the course of infection compared to wildtype controls (**Figure 8B** and **Supplementary Figure S5B**), suggesting that IL-17A signalling regulates both bodyweight and food intake. Next, we assessed whether adipocyte IL-17A signalling influenced iWAT mass during *T. brucei* infection. As with mice globally deficient for IL-17A/F (**Figure 5F**), infected *Adipoq^Cre^*x *Il17ra^fl/fl^* mice also experienced less iWAT wasting compared with C57BL/6 mice (**Figure 8C**). We also found that in contrast to naïve C57BL/6 mice, *Adipoq^Cre^* x *Il17ra^fl/fl^* mice had a higher frequency of small adipocytes, particularly in the 100-300 µm^2^ range (**Figure 8D and 8E**). Infected *Adipoq^Cre^*x *Il17ra^fl/fl^* mice also had higher frequencies of small adipocytes in the 51-200 µm^2^ range, compared with C57BL/6 mice (**Figure 8F**). The significantly higher frequency of smaller adipocytes in the *Adipoq^Cre^* x *Il17ra^fl/fl^* mice arise from an accumulation of interstitial preadipocytes, as shown in our single cell dataset (**Figure 7**), which are unable to differentiate to mature adipocytes. Together, this suggests that while the infected *Adipoq^Cre^*x *Il17ra^fl/fl^* mice retain more iWAT mass, the composition of the tissue is changed compared with C57BL/6 mice. To test this, we measured the expression of *Pi16* and *Dpp4*, which were exclusively upregulated in preadipocytes in our single cell dataset (**Supplementary Figure S5C**), as well as *Pparg* as a marker of mature adipocytes. Although we observed no significant increase in *Dpp4* or *Pi16* expression in the iWAT of infected C57BL/6 mice, both genes were significantly upregulated in the iWAT of infected *Adipoq^Cre^* x *Il17ra^fl/fl^* mice (**Figure 8G**). In addition, *Pparg* was upregulated in the iWAT of infected C57BL/6 mice, but unchanged in infected *Adipoq^Cre^* x *Il17ra^fl/fl^* mice (**Figure 8G**). Together, these data indicate that in the absence of functional IL-17 signalling on adipocytes, the *Pi16^+^ Dpp4^+^* interstitial preadipocytes are unable to complete their differentiation and/or development programme successfully towards *Pparg+* mature adipocytes (**Figure 8G**). Unexpectedly, we also found significantly higher number of parasites in the iWAT of *Adipoq^Cre^* x *Il17ra^fl/fl^* compared with C57BL/6 mice (**Figure 8H and 8I**), indicating that signalling through the adipocyte IL-17A receptor plays a central role in controlling local parasite numbers. These data demonstrate that IL-17 signalling is critical for promoting and/or supporting preadipocyte development towards mature adipocytes in the context of *T. brucei* infection, as well as impacting local parasite tissue burden.

**Figure 8.**
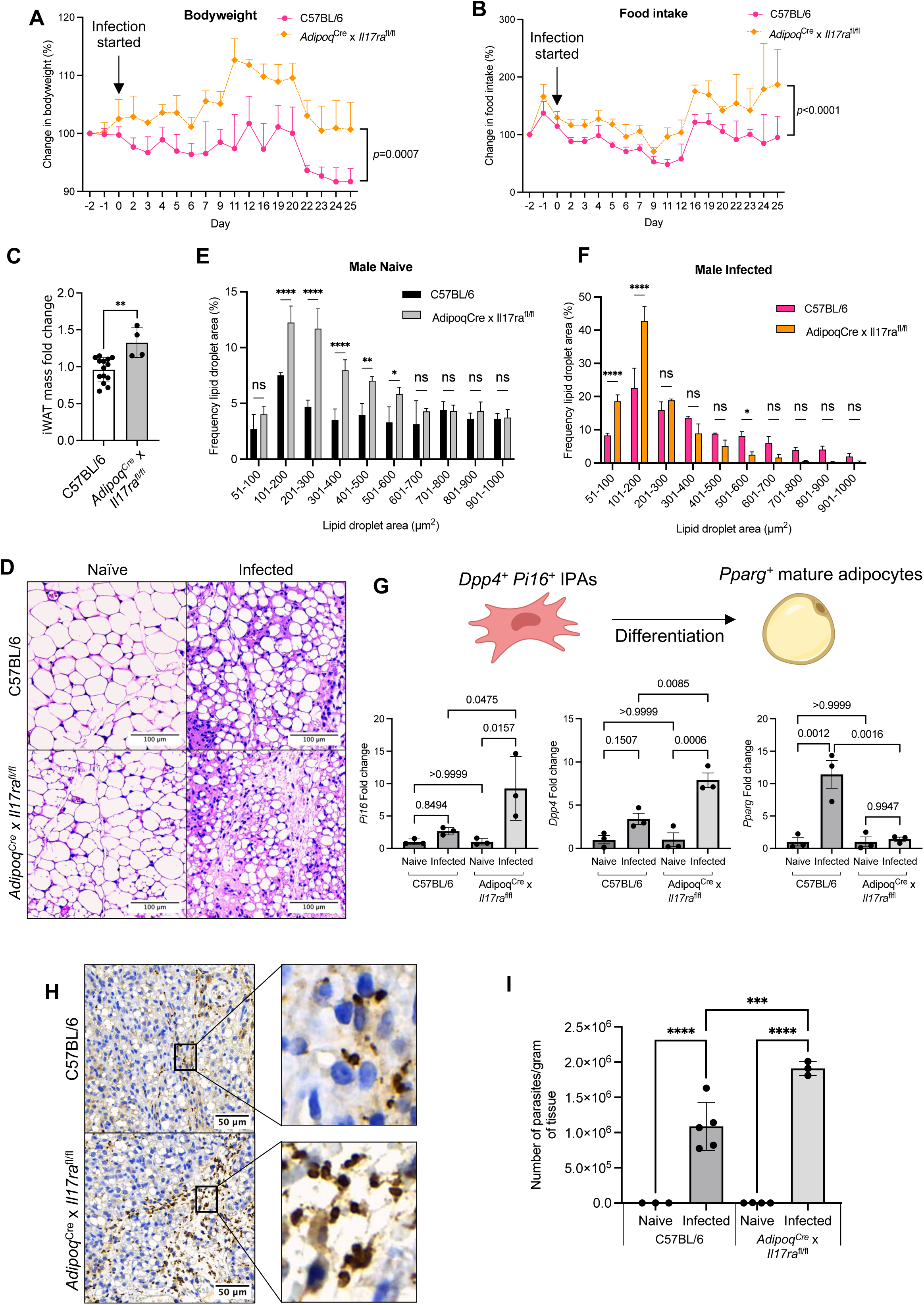
Adipocyte IL-17A signalling is critical for control of parasite numbers in the iWAT. (**A**) Percentage changes in body weight of male C57BL/6 or *Adipoq*^Cre^ x *Il17ra*^Flox^ mice. Percentage changes in food intake of singly housed male C57BL/6 or *Adipoq*^Cre^ x *Il17ra*^Flox^ mice. (**C**) iWAT mass at 25 days post-infection or in naïve mice. iWAT was dissected and weighed before normalising to body weight, to account for variation between biological replicates. Symbols indicate the number of biological replicates collected from two independent experiments. (**D**) Representative histological H&E staining of iWAT showing adipocyte lipid droplets and immune infiltrate. Analysis of lipid droplet area (µm^2^) in naïve (**E**) and infected (**F**) males. *n*=3-4 biological replicates per group. Lipid droplets were measured from 3 distinct areas in each image and then combined for each biological replicate. (**G**) RT-qPCR of *Pi16*, *Dpp4*, and *Pparg* from the iWAT. Cartoon illustrates gene expression by interstitial preadipocytes (IPAs) or mature adipocytes. (**H**) Histological analysis of the iWAT trypanosome colonisation. (**I**) Parasite density in the iWAT, which was measured by RT-qPCR of genomic DNA. For **A**, **B**, **E** and **F**, data were analysed using a two-way ANOVA with Sidak post-hoc testing. For **C**, data were analysed using a student’s t-test. For **G,** data were analysed using a one-way ANOVA with Tukey’s post-hoc testing. Biological replicates are indicated by symbols for histograms. For other plots, *n*=4 biological replicates/group. Data for all panels are expressed as mean ±SD. \**p*<0.05, \*\**p*<0.01, \*\*\**p*<0.001, \*\*\*\**p*<0.0001, ns = non-significant.

## Discussion

It is now clear that African trypanosomes establish dynamic interactions with several tissues, leading to the establishment of infectious foci important for disease transmission and pathogenesis. In this context, the adipose tissue has emerged as a critical site for survival, replication, and antigenic diversity, as recently shown^36^. In this study, we set out to determine the impact that African trypanosomes on the subcutaneous adipose tissue, which is adjacent to the skin and might be critical for forward transmission. Through various complementary analyses, we uncovered a previously unappreciated role of IL-17A/F in controlling tissue dynamics in response to *T. brucei* infection, acting on interstitial and/or committed preadipocytes to support adipocyte maturation and tissue replenishment under chronic inflammatory challenges. Notably, we also observed elevation of circulating IL-17A in HAT patients, suggesting that this cytokine plays a role in the human immune response to *T. brucei* infection. It is, therefore, tempting to speculate that IL-17A could also play a role in mediating the weight loss experienced by patients infected with *T. brucei*.

Previous studies have focused primarily on the use of a single sex during *T. brucei* infection, with a limited number of studies comparing sexes when measuring effects on reproductive organs^37, 38^. It is understood that males and females display multiple differential responses to infection, including differences in sickness behaviour^39^, weight loss^40^, and the immune response^41^. This aligns with the results presented here, where we observe differences in behaviour, weight loss and the immune response between male and female mice during infection. More specifically, weight loss in male mice was coupled with decreased iWAT mass, and was associated with a decrease in adipocyte lipid droplet size in male compared to female mice, potentially indicating that male mice utilise their lipid stores at a faster rate than females. Alternatively, this may be a result of females preferentially storing lipids in iWAT compared with males^42^, and so throughout infection they are able to store more fatty acids obtained from their diet than males, slowing down lipid droplet shrinkage. Due to this loss of lipid content, we hypothesised that circulating glycerol, which is a proxy for measuring adipocyte lipolysis^43^, would be elevated during infection. We found evidence suggesting that, during early infection, metabolic pathways such as glycolysis and lipolysis become more active, likely contributing to the changes in bodyweight and iWAT mass that we observed. However, we found that in both HAT patients and chronically infected mice circulating glycerol levels are diminished during trypanosomiasis, which may occur due to the extensive loss of adipocytes and subsequent decreases in lipolysis. Thus, it is tempting to speculate that since lipolysis is a critical regulator of multiple immune compartments, including macrophages^44^ and CD4^+^ T cells, that downregulation of this pathway will also impair the immune response to infection. Further work is required to elucidate if this also takes places in African trypanosomiasis.

To further understand the processes driving the adipocyte shrinkage that we observed during infection, we performed bulk transcriptomic analyses on the iWAT. This supported our findings that adipose tissue lipolysis is diminished during chronic infection, showing extensive downregulation of genes associated with mitochondrial metabolism and lipolysis in both males and female mice. This suppression of metabolic pathways has been observed in response to infection with other pathogens, such as *Mycobacterium tuberculosis*^46^. During *M. tuberculosis* infection of the lungs, transcripts encoding enzymes of oxidative phosphorylation and the tricarboxylic acid cycle are downregulated, and there is a shift to glycolysis^46^, which may support the inflammatory response to infection^47^. Furthermore, in diseases such as anorexia nervosa, prolonged weight loss and negative energy balance, lipolysis decreases, which is hypothesised to preserve energy that is needed to maintain processes essential to survival^48^. In the context of *T. brucei* infection, prolonged weight loss may result in the adipose tissue shutting down its main metabolic functions during chronic infection (namely lipolysis and mitochondrial metabolism) to preserve energy stores. Alternatively, it is also possible that the observed downregulation of metabolic transcripts in iWAT in response to infection could be caused by the loss of fully differentiated adipocytes, and therefore, underrepresentation of transcripts from these cells.

Previous studies of the adipose tissue immune response to *T. brucei* infection focused exclusively on the gWAT of male mice, and identified an expansion of macrophage, Teff cell, and B cell populations^37^. In agreement with this study, we also observed an expansion of macrophages and Teff cells. However, we also noted that there was a reduction in the proportion of B cells in the iWAT of male mice, and no changes in females, which could suggest that this is a tissue- and sex-specific response. We also observed an expansion of IL-17 producing T cells by CyTOF, which validated our transcriptomics dataset. This observation led us to question whether IL-17 plays a role in controlling iWAT tissue structure and/or function during *T. brucei* infection. Deficiency of IL-17A/F prevented weight loss in *T. brucei*-infected male mice compared to immunocompetent wild type male mice, but had no effect of bodyweight on female mice. When examining the tissue responses to infection at the single cell level, we identified that both *Dpp4^+^ Pi16^+^* interstitial and *Plin2^+^* committed preadipocytes significantly upregulate *Il17ra* in response to *T. brucei* infection, indicating that the capacity of these cells to sense IL-17 signalling is augmented during infection. These adipogenic precursor cells are critical for maintaining a normal supply of fully mature adipocytes under homeostatic conditions^32^, but their dynamics during infection remain poorly understood. Using a conditional deletion system to specifically delete *Il17ra* on adipocytes, we were able to identify a significant increase in the expression levels of *Dpp4* and *Pi16*, consistent with an accumulation of immature adipocytes. In addition to this, we found that adipocyte *Il17ra* deletion prevented upregulation of *Pparg*, further suggesting that these mice are less able to develop mature adipocytes during infection. Furthermore, at the histological level, this targeted deletion resulted in a higher proportion of small adipocytes in the Il17ra-deficient mice compared to the wildtype controls, under both naïve and infection conditions. This is in agreement with previous studies showing that deficiency of the adipocyte IL-17A receptor leads to reductions in iWAT lipid content under steady-state conditions^49^. In these previous studies, the reductions in lipid content resulted from the induction of thermogenesis in the iWAT, when adipocyte IL-17A signalling was inhibited, leading to increased energy substrate utilisation. Whilst there is evidence to suggest that non-shivering thermogenesis can be activated during infections such as influenza^50^, we did not find evidence of thermogenic activation during *T. brucei* infection, which may indicate that IL-17A signalling can also drive loss of iWAT mass by potentially impairing adipocyte maturation, which has been shown to occur *in vitro*^9^.

Unexpectedly, we found that unlike global deficiency of IL-17A/F, deficiency of the adipocyte IL-17A receptor diminished control of iWAT parasite burden, which raises the possibility that adipocytes are a key contributor to the immune response to *T. brucei* infection. This discrepancy may arise since the IL-17A receptor heterodimerises to the IL-17C receptor and, therefore, transduces signals from IL-17A, IL-17F, and IL-17C^51^, raising the possibility that IL-17C is critical for the control of local parasite numbers. Regardless, it remains to be determined whether loss of control of parasite numbers occurs because IL-17A receptor signalling is driving changes in lipolysis, or because IL-17A receptor signalling is altering the phenotype of adipocytes in a way that is influencing the local immune response. Supporting the former possibility, we recently used scRNAseq-based cell-cell communication analyses to show that adipocytes upregulate a range of factors, including *Cd40*, *Icam1*, *Jag1*, *Tnf*, and *Tnfsf18* in response to *T. brucei* infection, which could contribute to T cell activation^49^. Furthermore, in the same study we found that preadipocytes upregulate expression of inflammatory cytokines and antigen presentation molecules^17^. The inability to control local parasite numbers when adipocyte IL-17A receptor signalling is abrogated suggests that adipocytes are critical contributors to and coordinators of the local immune response to *T. brucei* infection, either by direct engagement with the immune system (e.g., *via* T cell activation), through metabolic fuelling (e.g., lipid mobilisation and release), or both. Recent work also found that when adipocytes can no longer engage lipolysis, there is a transient increase in the number of parasites in the gonadal adipose tissue early during infection^52^. The transient nature of this change could suggest that lipolysis is fuelling a specific arm of the immune response to infection that is necessary for parasite control at that time point. Although there is the possibility that lipolysis can limit parasite numbers due to lipotoxicity^52^, the parasites also upregulate genes associated with fatty acid oxidation when they reside in the adipose tissue^6^, potentially indicating that they are capable of adapting to the lipid-rich environment and may be resistant to lipotoxicity. The findings that we present here show that when local IL-17A receptor signalling is abrogated the architecture of the iWAT changes to contain more immature preadipocytes compared to wildtype controls, impacting the efficiency of the local immune response, and resulting in a higher parasite burden, once more placing adipocyte dynamics at the core of the immunological responses against *T. brucei* in the adipose tissue.

Based on all these observations, we propose a model whereby chronic *T. brucei* infection leads to an increase in circulating IL-17A/F in humans and animals, which act locally in the adipose tissue, by direct signalling through the adipocyte IL-17 receptor, and that this, in turn, acts to coordinate adipocyte maturation, local immune response, and efficient and timely parasite control. This study provides insights into the metabolic response of the subcutaneous adipose tissue during infection with *T. brucei* and how changes in cellular metabolism influence systemic metabolism. Future work employing lineage tracing studies are necessary to understand how adipose tissue dynamics are coupled to immunological responses in the context of chronic infections.

## Figure Legends

## Supplementary Figure Legends

**Supplementary Figure S1.** Spleen weights were normalised to bodyweight and the fold change in weight calculated. Each point represents a biological replicate. Data were analysed using one-way ANOVA with Tukey post-hoc testing. Data are expressed as mean ±SD. ****p<0.0001.

**Supplementary Figure S2.** Heatmaps, clustered by Euclidean distance, showing downregulation of lipolysis-related genes in male (**A**) and female (**B**) mice during chronic *T. brucei* infection

**Supplementary Figure S3**. (**A**) Number of parasites per mL of blood, measured using phase microscopy and the rapid “matching” method^53^. (**B**) Clinical scores of infected female mice. **(C)** Histological analysis of the iWAT and gWAT trypanosome colonisation, using HSP70 staining. (**D**) Parasite burden of iWAT and gWAT, which was measured by RT-qPCR of genomic DNA. (**E**) Percentage changes in body weight of female mice over the course of infection. (**F**) Percentage change in gross food intake. Each data point represents 2 cages (*n*=3-4 mice per cage). Spleen weights from males (**G**) and females (**H**) were normalised to bodyweight and the fold change in weight calculated. (**I**) iWAT mass at 25 days post-infection or in naïve mice. iWAT was dissected and weighed before normalising to body weight, to account for variation between biological replicates. (**J**) Representative histological H&E staining of iWAT showing adipocyte lipid droplets and immune infiltrate. (**K**) Analysis of lipid droplet area (µm^2^) in naïve and infected females. *N*=6 biological replicates per group, from two independent experiments. Lipid droplets were measured from 3 distinct areas in each image and then combined for each biological replicate. Time series data were analysed using two-way repeated measures ANOVA with Sidak post-hoc testing. **D**, **G**, and **H** were analysed using a one-way ANOVA with Tukey’s post-hoc testing. **I** was analysed using a student’s t-test. Data for all panels are expressed as mean ±SD. \**p*<0.05, \*\*\*\**p*<0.0001, ns = non-significant.

**Supplementary Figure S4**. Dot plot showing expression of genes involved in glycolysis and the TCA cycle (**A**) or the pentose phosphate pathway (**B**). In the key, dots represent the percentage of cells that express a given marker, and the colour intensity represent the level of expression. Single cell data are comprised of one technical replicate per condition, each containing pooled cells from the iWAT of 5 male mice replicate.

**Supplementary Figure S5**. (**A**) Three-way comparison of the bodyweight of single-housed infected C57BL/6, *Il17af*-/- and *Adipoq*^Cre^ x *Il17ra*^Flox^ mice. (**B**) Three-way comparison of the food intake of single-housed infected C57BL/6, *Il17af*-/- and *Adipoq*^Cre^ x *Il17ra*^Flox^ mice. (**C**)) Dot plot from scRNAseq showing expression of *Dpp4* and *Pi16* across all cell populations. Time series data were analysed using two-way repeated measures ANOVA with Sidak post-hoc testing. Data for all panels are expressed as mean ±SD.

## Materials & Methods

### Ethics statement

Male and female human serum samples used for IL-17A measurements were collected in Guinea as part of the National Control Program. Participants were informed of the study objectives in their own language and signed a written consent form. Participants comprised six males and four females, all over the age of 18 years. Approval for this study was obtained from the Comité Consultative de Déontologie et d’Ethique (CCDE) of the Institut de Recherche pour le Développement (approval number 1–22/04/2013). Due to limited sample numbers, we did not stratify data by sex. Human serum samples used for glycerol measurements were collected as part of the TrypanoGEN Biobank^54^, with ethical approval from the Democratic Republic of Congo National Ministry of Public Health (approval number 1/2013). Samples were used from different regions due to limitations in sample availability. Ethical approval to use all human samples outlined in this study was given by the University of Glasgow (approval number 200120043). All animal experiments were approved by the University of Glasgow Ethical Review Committee and performed in accordance with the home office guidelines, UK Animals (Scientific Procedures) Act, 1986 and EU directive 2010/63/EU. All experiments were conducted under SAPO regulations and UK Home Office project licence number PC8C3B25C and PP4863348 to Dr Jean Rodgers and Professor Annette MacLeod, respectively.

### Mouse generation and infections with *Trypanosoma brucei*

Eight to ten-week old Adipoq-cre (JAX, stock 028020) or Il17ratm2.1Koll/J (JAX, stock 031000) were purchased from The Jackson Laboratory. These mice were crossed to generate Adipoq^Cre^ x Il17ra^Flox^ mice, with an adipocyte specific deletion of the IL-17A receptor. Six to eight weeks old male or female C57BL/6J (JAX, stock 000664), *Il17af^-/-^* (JAX, stock 034140), or IL-17A GFP reporter mice (JAX, stock 018472) were also purchased. At >10 weeks old, mice were randomly allocated to control or treatment groups by animal unit technical staff. Mice were then inoculated by intra-peritoneal injection with ∼2 x 10^3^ of the *T. brucei brucei* Antat 1.1^E^ parasite strain^55^. Parasitaemia was monitored by regular sampling from tail venesection and examined using phase microscopy and the rapid “matching” method^53^. Uninfected mice of the same strain, sex and age served as uninfected controls. All mice were fed *ad libitum* and kept on a 12 h light–dark cycle. All *in vivo* experiments were concluded at 25 days post-infection, to model chronic infection in humans.

### RNA Purification and Bulk RNA sequencing

iWAT was harvested and stored in TRIzol™ (Invitrogen). Total RNA was then purified using an RNeasy Kit (Qiagen) as per the manufacturer’s recommendations. The RNA was purified in 30 µL of nuclease-free water (Qiagen), and RNA concentration measured on a NanoDrop™ 2000 (Thermo Fisher Scientific). Samples were shipped to Novogene (Cambridge, UK) to undergo quality control, library preparation and sequencing. RNA integrity was assessed using an RNA Nano 6000 Assay Kit (Agilent Technologies) with a Bioanalyzer 2100 (Agilent Technologies), as per the manufacturer’s instructions. Samples with an RNA integrity number (RIN) of >6.0 were qualified for RNA sequencing.

#### Library Preparation

Library preparation was performed by Novogene (Cambridge, UK). Messenger RNA (mRNA) was purified from total RNA using poly-T oligo-attached magnetic beads. Fragmentation was carried out using divalent cations under elevated temperature in a First Strand Synthesis Reaction Buffer (5X). First strand cDNA was synthesized using random hexamer primers and M-MuLV Reverse Transcriptase (RNase H-). Second strand cDNA synthesis was then performed using DNA Polymerase I and RNase H. Remaining overhangs were converted to blunt ends *via* exonuclease/polymerase activity. Following adenylation of 3’ ends of DNA fragments, adaptors with hairpin loop structures were ligated. To select cDNA fragments of 370∼420 bp in length, library fragments were purified using AMPure XP beads (Beckman Coulter), as per the manufacturer’s instructions. PCR was then performed using Phusion High-Fidelity DNA polymerase, Universal PCR primers, and Index (X) primers. Finally, PCR products were purified (AMPure XP system) using AMPure XP beads (Beckman Coulter), as per the manufacturer’s instructions, and library quality was assessed using a Bioanalyzer 2100 (Agilent Technologies).

#### Sequencing and data analysis

Clustering of the index-coded samples was performed on a cBot Cluster Generation System using a TruSeq PE Cluster Kit v3-cBot-HS (Illumia) according to the manufacturer’s instructions. After cluster generation, libraries were sequenced on an Illumina Novaseq platform and 150 bp paired-end reads were generated. Raw reads in fastq format were processed through proprietary Perl scripts developed by Novogene (Cambridge, UK). Clean reads were obtained by removing reads containing adapters, poly-N, or low-quality reads from raw data. Concurrently, the Q20, Q30 and GC content of the clean data was calculated. Genome and genome annotation files (Genome Reference Consortium Mouse Build; GRCm39) were downloaded. An index of the reference genome was built using Hisat2 v2.0.5 and paired-end clean reads were aligned to the reference genome using Hisat2 v2.0.5. The featureCounts (v. 1.5.0-p3) package was used to count read numbers mapped to each gene, before calculating the Fragments Per Kilobase of transcript sequence per Millions base pairs (FPKM) of each gene using the length of the gene and reads count mapped to this gene. Differential expression analysis was performed using the DESeq2 R package (v. 1.20.0). DESeq2 The resulting *p*-values were adjusted using the Benjamini and Hochberg approach to control false discovery rate. Genes with an adjusted *p*-value of <0.05 were assigned as differentially expressed. Pathway enrichment analysis of differentially expressed genes was performed using the ShinyGO (v. 0.76) package, mapping genes to the Kyoto Encyclopaedia of of Genes and Genomes (KEGG) database. KEGG terms with an adjusted *p*-value <0.05 were considered significantly enriched. Heatmaps were generated using the pheatmap (Version 1.0.12) and Tidyverse packages in R (Version 4.2.1). Samples were clustered by Euclidean distance.

### Adipose tissue processing and preparation of single cell suspension for single-cell RNA sequencing

Infected animals and naïve controls were anesthetized with isoflurane at 7 days post-infection and perfused transcardially with 25-30 ml of ice-cold 1X PBS containing 0.025% (wt/vol) EDTA, and both iWAT pads were excised, and the inguinal lymph node removed. For each condition, fat pads were pooled from 5 mice, resulting in a single technical replicate per condition. The iWAT was dissociated using an Adipose Tissue Dissociation Kit, mouse and rat (Miltenyi Biotec) with a gentleMACS™ Octo Dissociator with Heaters (Miltenyi Biotec) as per the manufacturer’s recommendations. Digested tissue was then passed through a 70 µm and then a 40 µm nylon mesh filter, which were washed with DMEM. The suspension was then centrifuged at 400 x g at 4°C for 5 minutes to isolate the stromal vascular fraction and remove adipocytes. Finally, cells were passed through a MACS dead cell removal kit (Miltenyi Biotec) and diluted to ∼1,000 cells/μl in 200μl HBSS 0.04% BSA and kept on ice until single-cell capture using the 10X Chromium platform. The single cell suspensions were loaded onto independent single channels of a Chromium Controller (10X Genomics) single-cell platform. Single cells were loaded for capture using 10X Chromium NextGEM Single cell 3 Reagent kit v3.1 (10X Genomics). Following capture and lysis, complementary DNA was synthesized and amplified (12 cycles) as per the manufacturer’s protocol (10X Genomics). The final library preparation was carried out as recommended by the manufacturer with a total of 14 cycles of amplification. The amplified cDNA was used as input to construct an Illumina sequencing library and sequenced on a Novaseq 6000 sequencers by Glasgow Polyomics.

### Read mapping, data processing, and integration

For FASTQ generation and alignments, Illumina basecall files (*.bcl) were converted to FASTQs using bcl2fastq. Gene counts were generated using Cellranger v.6.0.0 pipeline against a combined Mus musculus (mm10) and Trypanosoma brucei (TREU927) transcriptome reference. After alignment, reads were grouped based on barcode sequences and demultiplexed using the Unique Molecular Identifiers (UMIs). The mouse-specific digital expression matrices (DEMs) from all six samples were processed using the R (v4.2.1) package Seurat v4.1.017. Additional packages used for scRNAseq analysis included dplyr v1.0.7 18, RColorBrewer v1.1.2 (http://colorbrewer.org), ggplot v3.3.5 19, and sctransform v0.3.3 20. We initially captured 50,640 cells mapping specifically against the *M. musculus* genome across all conditions and biological replicates, with an average of 20,209 reads/cell and a median of 1,302 genes/cell (**Supplementary Table 3**). The number of UMIs was then counted for each gene in each cell to generate the digital expression matrix (DEM). Low quality cells were identified according to the following criteria and filtered out: i) nFeature > 100 or <5,000, ii) nCounts > 100 or <20,000, iii) > 30% reads mapping to mitochondrial genes, and iv) > 40% reads mapping to ribosomal genes, v) genes detected < 3 cells. After applying this cut-off, we obtained a total of 46,546 high quality mouse-specific cells with a median of 1,296 genes/cell (**Supplementary Table 3**). High-quality cells were then normalised using the SCTransform function, regressing out for total UMI and genes counts, cell cycle genes, a nd highly variable genes identified by both Seurat and Scater packages, followed by data integration using IntegrateData and FindIntegrationAnchors. For this, the number of principal components were chosen using the elbow point in a plot ranking principal components and the percentage of variance explained.

### Cluster analysis, marker gene identification, and subclustering

The integrated dataset was then analysed using RunUMAP (10 dimensions), followed by FindNeighbors (10 dimensions, reduction = “pca”) and FindClusters (resolution = 0.4). With this approach, we identified a total of 16 cell clusters The cluster markers were then found using the FindAllMarkers function (logfc.threshold = 0.25, assay = “RNA”). To identify cell identity confidently, we employed a supervised approach. This required the manual inspection of the marker gene list followed by and assignment of cell identity based on the expression of putative marker genes expressed in the unidentified clusters. To increase the resolution of our clusters to help resolve potential mixed cell populations embedded within a single cluster, we subset preadipocytes and T cells and analysed them separately using the same functions described above. In all cases, upon subsetting, the resulting objects were reprocessed using the functions FindVariableFeatures, ScaleData, RunUMAP, FindNeighbors, and FindClusters with default parameters. The number of dimensions used in each case varied depending on the cell type being analysed but ranged between 5 and 10 dimensions. Cell type-level differential expression analysis between experimental conditions was conducted using the FindMarkers function (min.pct = 0.25, test.use = Wilcox) and (DefaultAssay = “SCT”).

### DNA Purification

Tissues were harvested from mice and snap frozen. Tissue was digested using a DNeasy Blood and Tissue kit (Qiagen), before purifying DNA as per the manufacturer’s instructions. DNA was eluted in 100 µL of EB buffer (Qiagen).

### Tissue Parasite Burden Quantification

To quantify *T. brucei* parasites in tissue, we amplified 18S ribosomal DNA genes from the gDNA of a known mass of tissue, using qRT-PCR Brilliant II Probe Master Mix (Agilent Technologies) with a TaqMan™ TAMRA Probe system (Applied Biosystems). Primer sequences were specific to *T. brucei* 18S ribosomal DNA (**Table 1**). The cycling conditions used for qRT-PCR are outlined in **Table 2**. Generated data was converted to parasite copy number using a standard curve.

**Table 1.**
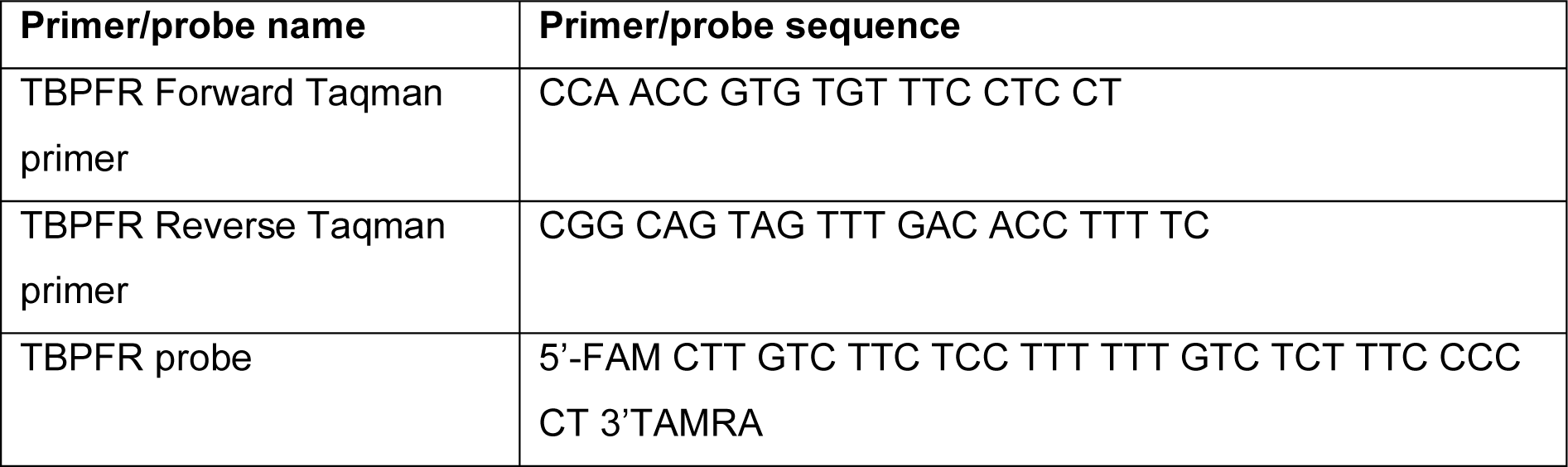
Primer and probe sequences for tissue parasite burden quantification

**Table 2.** Thermal cycling conditions for tissue parasite burden quantification

| Step | Temperature (°C) | Time | Number of cycles |
| --- | --- | --- | --- |
| 1 | 95 | 10 minutes | 1 |
| 2 | 95 | 15 seconds | 45 |
| 3 | 60 | 1 minutes |  |
| 4 | 72 | 1 seconds |  |

### Real-time quantitative PCR

Following RNA purification, cDNA was synthesised, and the RT-qPCR master mix prepared using a Luna Universal One-Step RT-qPCR Kit (New England Biolabs), as per the manufacturer’s instructions, with the primers outlined in **Table 3**. The RT-qPCR was run on a Roche LightCycler 480 (Roche Diagnostics Ltd) using the conditions outlined in **Table 4**.

**Table 3.**
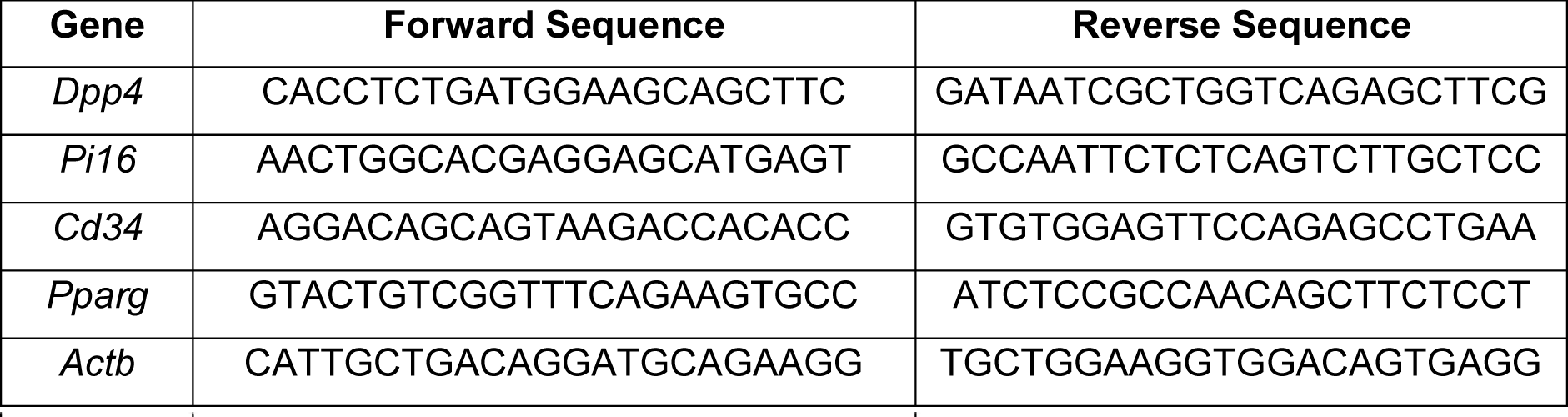
Primer sequences for RT-qPCR

**Table 4.**
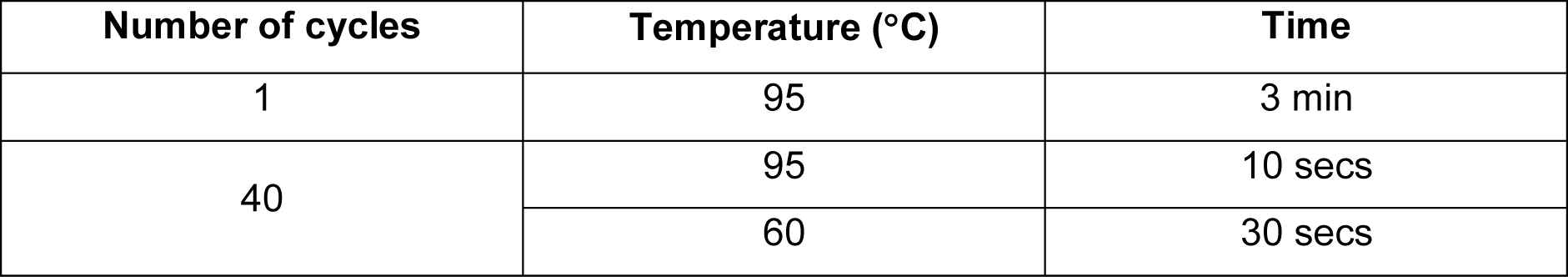
Cycling conditions for RT-qPCR

### Histological Analyses

Tissues were placed into 4% paraformaldehyde (PFA) and fixed overnight at room temperature. PFA-fixed tissues were then embedded in paraffin, sectioned, and stained by the Veterinary Diagnostic Services facility (University of Glasgow, UK). Sections were Haematoxylin and Eosin (H&E) stained for lipid droplet measurement analysis, or 3’-diaminobenzidine (DAB) stained for heat-shock protein 70 (HSP70) to detect *T. brucei* parasites. The HSP70 antibody was a kind gift from Professor James D. Bangs. Slide imaging was performed by the Veterinary Diagnostic Services facility (University of Glasgow, UK) using an EasyScan Infinity slide scanner (Motic) at 20X magnification. To determine lipid droplet sizes in adipose tissue, images were first opened in QuPath (v. 0.3.2)^56^, before selecting regions and exporting to Fiji^57^. In Fiji, images were converted to 16-bit format, and we used the Adiposoft plugin to quantify lipid droplet area within different sections.

### Mass cytometry sample processing

Adipose tissue was dissected out and transferred to PBS, before dissociating using an Adipose Tissue Dissociation Kit for Mouse and Rat (Miltenyi Biotec), using a gentleMACS™ Octo Dissociator with Heaters (Miltenyi Biotec), as per the manufacturer’s recommendations. After the final recommended centrifugation, the pellet (containing the immune cells) was resuspended in Dubecco’s Modified Eagle Medium (DMEM) to a concentration of 1 x 10^6^ cells/mL. Cells were activated for 3 h in a round-bottom 96-well plate using Cell Activation Cocktail (with Brefeldin A) (BioLegend). Plates were then centrifuged at 300 x g for 5 min and the pellets resuspended in 50 µL of Cell-ID™ Cisplatin-195Pt viability reagent (Standard BioTools), and incubated at room temperature for 2 min. Cells were washed twice in Maxpar® Cell Staining Buffer (Standard BioTools), and centrifuged at 300 x g at room temperature for 5 min. The CD16/CD32 receptors were then blocked by incubating with a 1/50 dilution of TruStain FcX™ (BioLegend) in PBS at room temperature for 15 min. An antibody cocktail was prepared (**Table 5**) from the Maxpar® Mouse Sp/LN Phenotyping Panel Kit (Standard BioTools) with additional antibodies against IL-17A, IFNγ, TCRgd, and CD27 included (**Table 5**). Cells were incubated with antibodies (**Table 5**) for 60 min, on ice before washing 3 times in Maxpar® Cell Staining Buffer (Standard BioTools) as previously.

**Table 5.** Antibodies for mass cytometry

| Surface Antibodies |  |  |  |  |
| --- | --- | --- | --- | --- |
| Antibody | Metal Conjugate | Clone | Concentration | Cat. No. |
| Ly6G/C [Gr1] | 141Pr | RB6-8C5 | 1/100 | 201306 |
| CD11c | 142Nd | N418 | 1/100 |  |
| CD69 | 145Nd | H1.2F3 | 1/100 |  |
| CD45 | 147Sm | 30-F11 | 1/200 |  |
| CD11b | 148Nd | M1/70 | 1/100 |  |
| CD19 | 149Sm | 6D5 | 1/100 |  |
| CD3e | 152Sm | 145-2C11 | 1/100 |  |
| TCRβ | 169Tm | H57-597 | 1/100 |  |
| CD44 | 171Yb | IM7 | 1/100 |  |
| CD4 | 172Yb | RM4-5 | 1/100 |  |
| Intracellular Antibodies |  |  |  |  |
| IL-17A | 174Yb | TC11-18H10.1 | 1/100 | 3174002C |
| IFNγ | 165Ho | XMG1.2 | 1/100 | 3165003B |

Following staining, cells were fixed in 2% paraformaldehyde (PFA) overnight at 4 °C. Cells were then washed twice with 1 x eBioscience™ Permeabilization Buffer (Invitrogen) at 800 x g at room temperature for 5 min. The pellets were resuspended in intracellular antibody cocktail (**Table 5**) and incubated at room temperature for 45 min. Cells were washed 3 times in Maxpar® Cell Staining Buffer (Standard BioTools) at 800 x g. The cells were then resuspended in 4% PFA at room temperature for 15 min, before collecting the cells at 800 x g and resuspending in Cell-ID™ Intercalator-Ir (Standard BioTools). Finally, the cells were barcoded by transferring the stained cells to a fresh tube containing 2 µL of palladium barcode from the Cell-ID™ 20-Plex Pd Barcoding Kit (Standard BioTools). Cells were then frozen in a freezing solution (90% FBS and 10% DMSO), before shipping to the Flow Cytometry Core Facility at the University of Manchester for data acquisition.

### Flow cytometric analyses

Infected animals and naïve controls were anesthetized with isoflurane and perfused transcardially with 25-30 ml of ice-cold 1X PBS containing 0.025% (wt/vol) EDTA. iWAT pads were excised and the inguinal lymph node was removed. The iWAT was dissociated using an Adipose Tissue Dissociation Kit, mouse and rat (Miltenyi Biotec) with a gentleMACS™ Octo Dissociator with Heaters (Miltenyi Biotec) as per the manufacturer’s recommendations. Digested tissue was then passed through a 70 µm and then a 40 µm nylon mesh filter, which were washed with DMEM. The suspension was then centrifuged at 400 x g at 4°C for 5 minutes to isolate the immune cell fraction and remove adipocytes. The resulting suspension was seeded on a 96-well plate and stimulated with 1X Cell Activation Cocktail containing phorbol 12-myristate 13-acetate (PMA), Ionomycin, and Brefeldin A (BioLegend) for 3 h at 37 °C and 5% CO_2_. For flow cytometry analysis, single cell suspensions were resuspended in ice-cold FACS buffer (2 mM EDTA, 5 U/ml DNAse I, 25 mM HEPES and 2.5% foetal calf serum (FCS) in 1X PBS) and stained for extracellular markers at 1:400 dilution (**Table 6**). Samples were run on a flow cytometer LSRFortessa (BD Biosciences) and analysed using FlowJo software version 10 (Treestar).

**Table 6.**
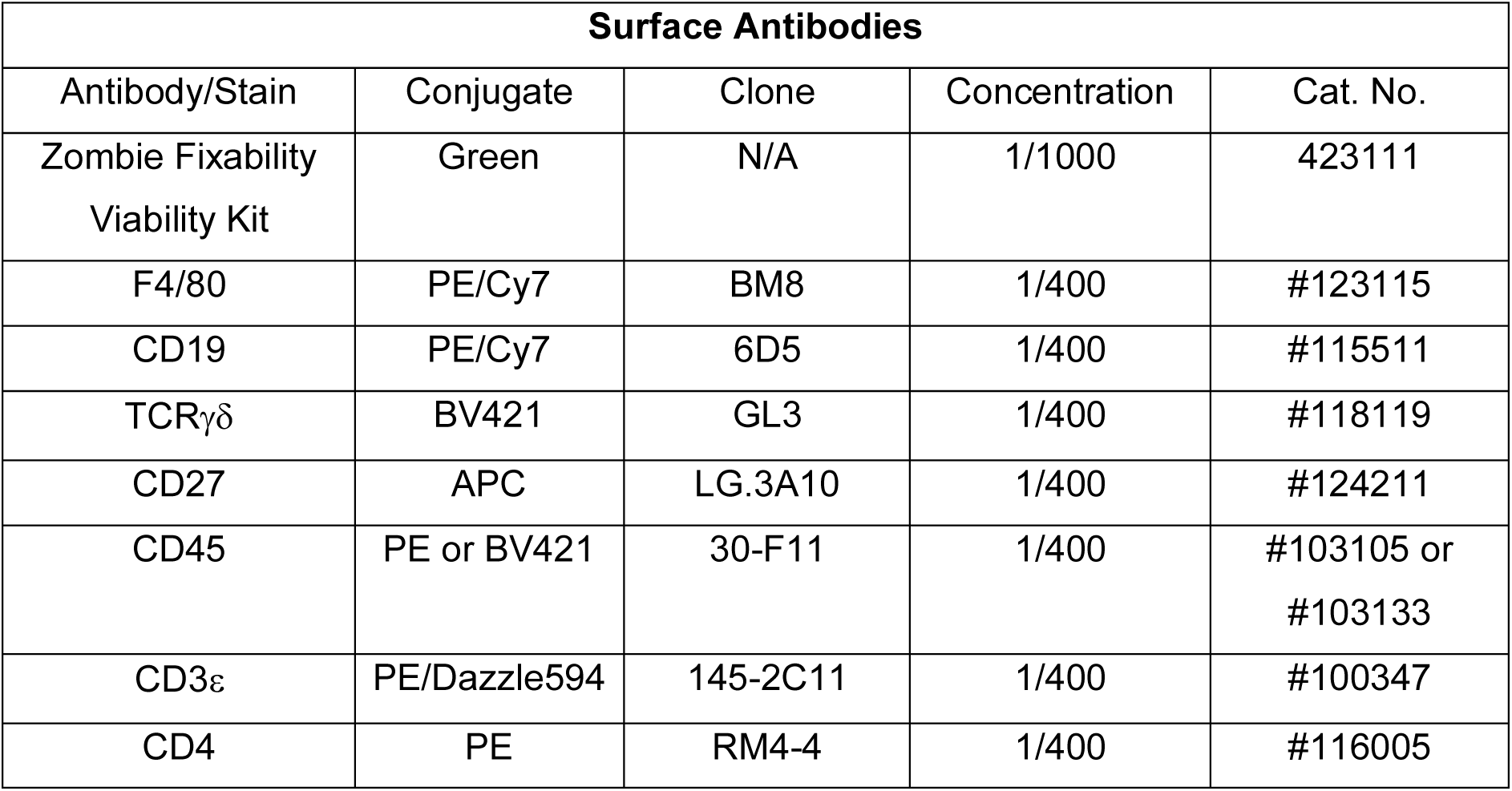
Antibodies for flow cytometry

| Surface Antibodies |  |  |  |  |
| --- | --- | --- | --- | --- |
| Antibody/Stain | Conjugate | Clone | Concentration | Cat. No. |
| Zombie Fixability Viability Kit | Green | N/A | 1/1000 | 423111 |
| F4/80 | PE/Cy7 | BM8 | 1/400 | #123115 |
| CD19 | PE/Cy7 | 6D5 | 1/400 | #115511 |
| TCR$\gamma\delta$ | BV421 | GL3 | 1/400 | #118119 |
| CD27 | APC | LG.3A10 | 1/400 | #124211 |
| CD45 | PE or BV421 | 30-F11 | 1/400 | #103105 or #103133 |
| CD3ε | PE/Dazzle594 | 145-2C11 | 1/400 | #100347 |
| CD4 | PE | RM4-4 | 1/400 | #116005 |

### Quantification of cytokine titres

To measure cytokine titres in murine serum samples we used a U-PLEX Biomarker kit (Meso Scale Discovery), as per the manufacturer’s instructions. Samples were analysed using a MESO QuickPlex SQ 120 (Meso Scale Discovery). IL-17A titres in human serum samples were quantified using a multiplex cytokine panel (Bio-Plex Pro Human Cytokine Assay, BioRad) and a LuminexCorp Luminex 100 machine as per the manufacturer’s instructions.

### Statistical analyses

All statistical analyses were performed using Graph Prism Version 8.0 for Windows or macOS, GraphPad Software (La Jolla California USA). Normality of data distribution was measured using the Shapiro-Wilks test. Where indicated, data were analysed by unpaired Student’s t-test, Mann-Whitney test, one-way analysis of variance (ANOVA) or two-way ANOVA. Data were considered to be significant where p <0.05.

## Data Availability

The GEO accession number for raw bulk transcriptomic sequencing and processed data reported in this paper is GSE210600. The GEO accession number for the raw scRNAseq and processed data reported in this paper is GSE233312. The scripts used to generate single cell data in this study are available at Zenodo (10.5281/zenodo.7966849).

## Acknowledgements

We firstly thank the TrypanoGEN Network for providing serum samples from patients. We also thank Jean Rodgers for the use of her project licence for performing animal work. We thank the Histology Research Service at Veterinary Diagnostic Services, School of Veterinary Medicine, University of Glasgow. We also thank Nicola Munro, Scott McCall and Catrina Boyd at the Veterinary Research Facility (University of Glasgow) for maintaining optimal husbandry conditions and comfort for the animals used in this study. We thank Julie Galbraith and Pawel Herzyk (Glasgow Polyomics, University of Glasgow) for their support with single cell library preparation and sequencing. Finally, the authors would like to thank the Flow Cytometry Core Facility, University of Manchester, UK, for mass cytometry sample acquisition. This work was funded in part by a Wellcome Trust Institutional Strategic Support Fund award [316917-01], a Society for Endocrinology Early Career Grant [316705/0], and a Wellcome Centre for Integrative Parasitology FutureScope grant [174811-23] to MCS. This work was also funded in part by a Wellcome Trust Senior Research Fellowship [209511/Z/17/Z] awarded to AML. JFQ is funded by a Sir Henry Wellcome postdoctoral fellowship (221640/Z/20/Z to JFQ). PC is funded by Wellcome Centre for Integrative Parasitology FutureScope grant to JFQ [104111/Z/14/Z]. GPW is funded by an MRC grant [MR/S009779/1]. CB is funded by an MRC grant [MR/W018497/1]. SK is funded by the National Institute of Diabetes and Digestive and Kidney Diseases and the Howard Hughes Medical Institute. The authors declare that the research was conducted in the absence of any commercial or financial relationships that could be construed as a potential conflict of interest.

