## Supplementary figures and images for "IL-17 signalling is critical for controlling subcutaneous adipose tissue dynamics and parasite burden during chronic Trypanosoma brucei infection"

### Figure S1

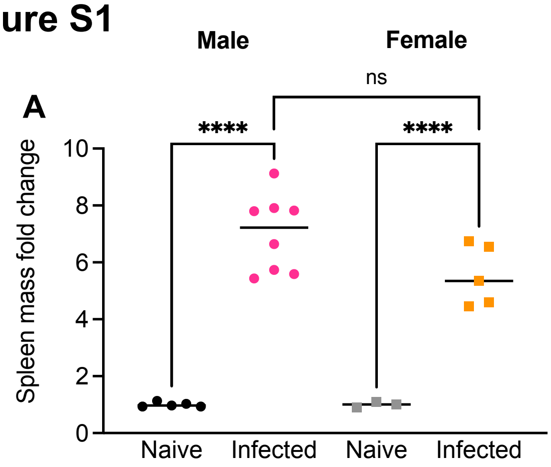

### Figure S2

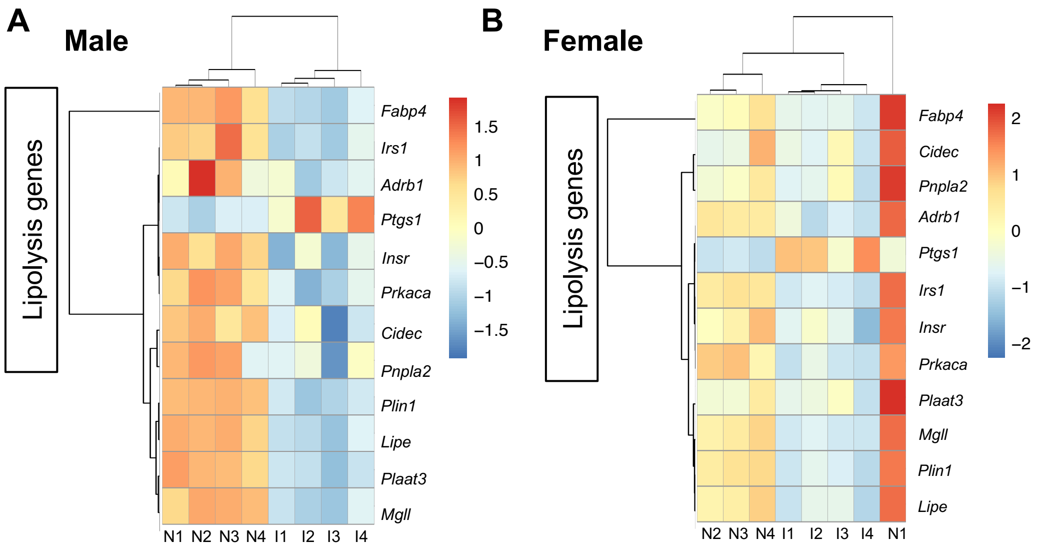

### Figure S3

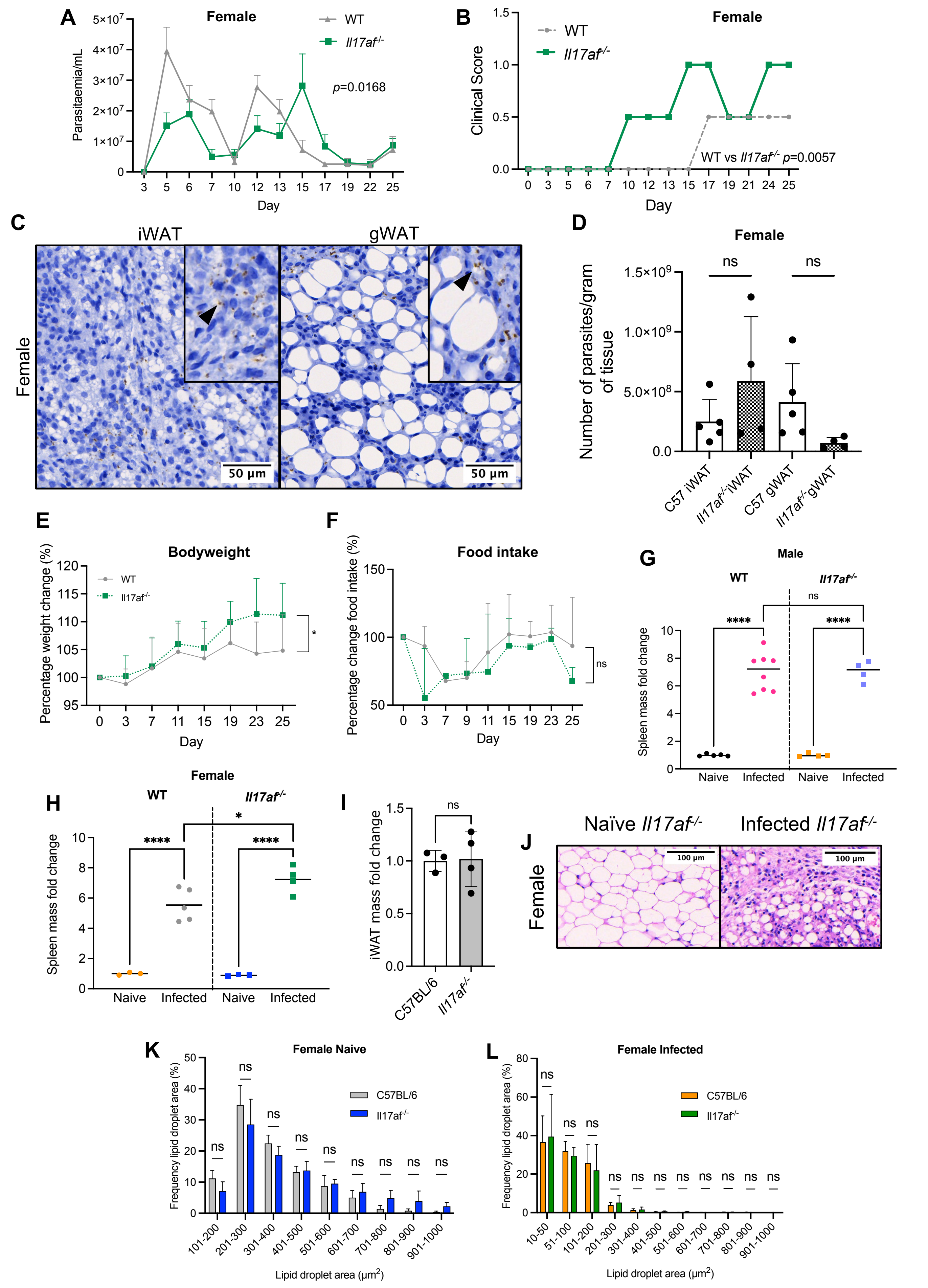

### Figure S4

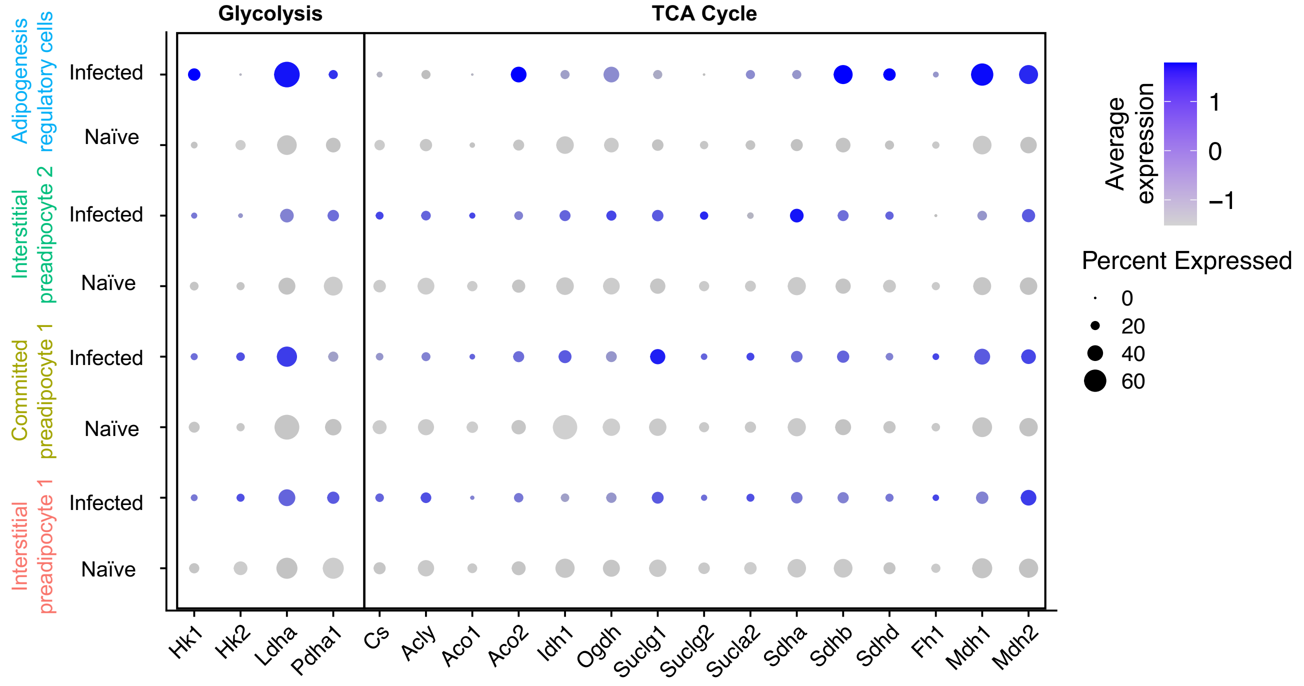

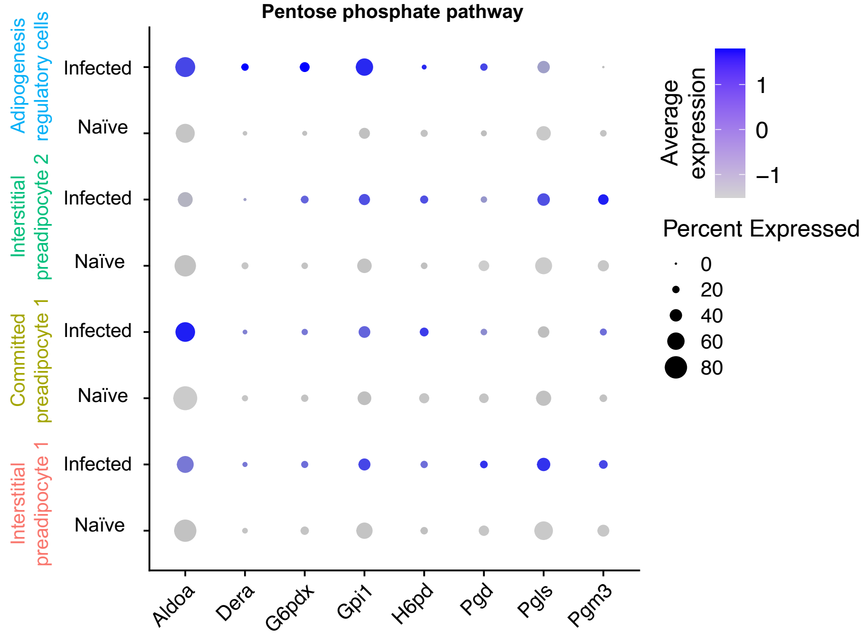

### Figure S5

#### Figure S5

A

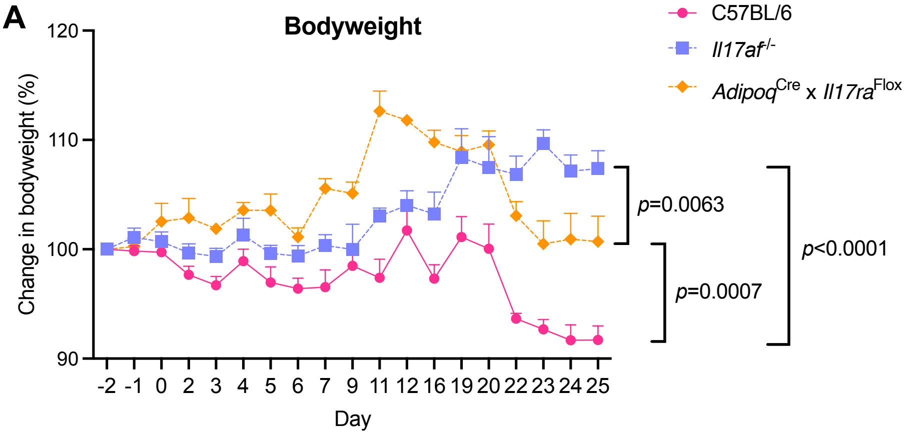


B

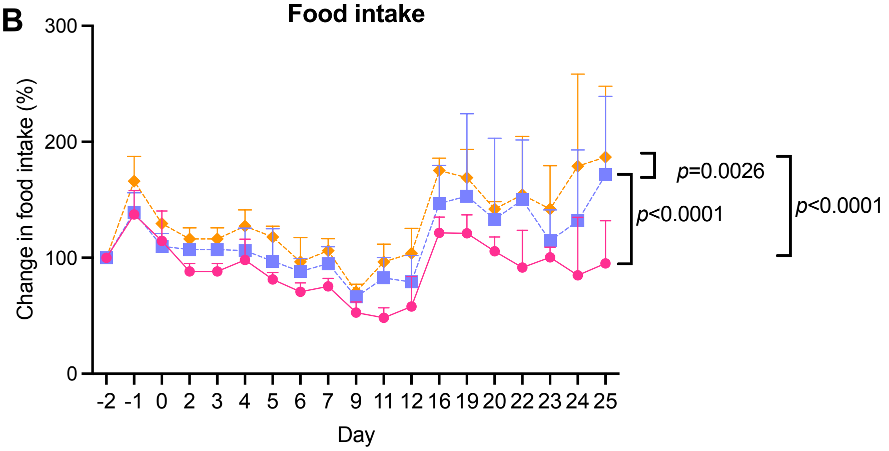


C

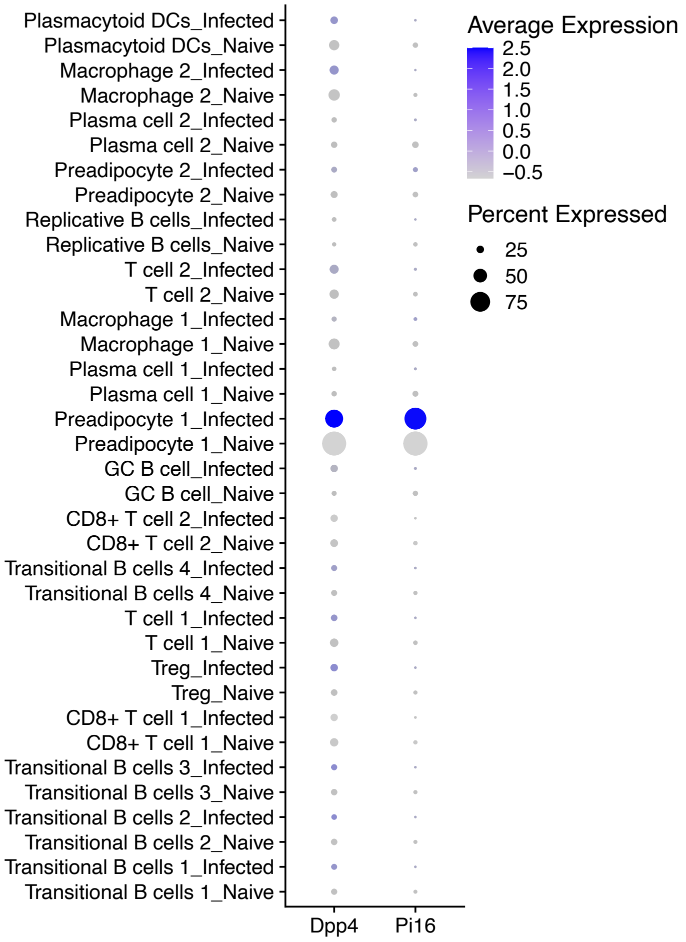
